# UniST: A Unified Computational Framework for 3D Spatial Transcriptomics Reconstruction

**DOI:** 10.64898/2026.03.12.711362

**Authors:** Lan Shui, Yunhe Liu, Idania Carolina Lubo Julio, Jean R Clemenceau, Xen Ping Hoi, Yibo Dai, Wei Lu, Jimin Min, Khaja Khan, Bailey Roemer, Mei Jiang, Rebecca Elaine Waters, Karen Colbert, Anirban Maitra, Max Wintermark, Ying Yuan, Keith Syson Chan, Tae Hyun Hwang, Paul F Mansfield, Jeremy Davis, Luisa M Solis Soto, Linghua Wang, Liang Li, Ziyi Li

## Abstract

Spatial transcriptomics (ST) enables the measurement of gene expression in its native spatial context, yet most ST datasets are acquired as two-dimensional (2D) sections. Consequently, the underlying three-dimensional (3D) organization of tissues is only partially observed, and 3D ST data generated from serial sections are typically sparse and heterogeneous, with substantial tissue loss and missing measurements. These limitations pose major analytical challenges for reconstructing coherent 3D tissue architecture, rather than issues of experimental scalability alone. Here, we present UniST, a unified generative artificial intelligence (AI) framework designed to computationally reconstruct dense and continuous 3D ST landscapes from sparse serial sections, without altering the underlying experimental ST technologies. UniST integrates three complementary modules: kernel point convolution with cross-attention layers for point cloud upsampling, optical flow-based interpolation for continuous slice reconstruction, and a graph autoencoder with implicit neural representations for gene expression imputation. Together, these components densify sparse slices, resolve discontinuities, and map spatial coordinates to high-dimensional transcriptomics. Across multiple ST platforms and tissue contexts, UniST accurately restored structural continuity and biologically meaningful expression patterns. In a mouse embryo dataset, UniST reconstructed a dense 3D heart architecture from sparsely sampled slices. In two 3D human cancer tissues, UniST recovered critical spatial features, including tumor-immune boundaries and tertiary lymphoid structures, that were fragmented in the original data. By providing a generalizable computational solution that complements existing ST acquisition protocols, UniST facilitates cost-efficient and scalable reconstruction of 3D ST landscapes, enabling more faithful investigation of tissue organization and disease biology.

## Introduction

Spatial transcriptomics (ST) profiles gene expression *in situ* and has become a fundamental tool for studying tissue organization, cellular heterogeneity, and microenvironmental structure across diverse biological contexts [1–5]. However, most ST datasets are acquired as two-dimensional (2D) tissue sections, providing only a limited view of the underlying three-dimensional (3D) tissue architecture. As a result, spatial relationship along the *z*−axis are incompletely captured, motivating growing interest in extending ST analyses to 3D context [6–8].

In practice, current efforts to extend ST from 2D to 3D largely rely on serial section-based strategies [9–12]. While these approaches enable initial exploration of tissue-scale organization, they also introduce fundamental data limitations. Individual tissue sections typically measure only 5∼10μm in thickness, thereby reconstructing large tissue volumes requires hundreds or thousands of consecutive slices, making comprehensive profiling experimentally burdensome and prohibitively expensive. Consequently, many studies selectively profile only a sparse subset of sections to manage cost, resulting in discontinuous *z*−axis coverage, missing measurements, and incomplete 3D representations [9, 13].

Beyond experimental burden, serial section-based 3D ST datasets are characterized by substantial missingness and heterogeneity. Tissue sections may be damaged or lost during processing, alignment across slices is imperfect, and sparsely sampled measurements along the *z*−axis yield fragmented “2.5D” approximations rather than continuous 3D tissue structures. Together, these challenges fundamentally limit downstream spatial analysis and biological interpretation, highlighting the need for computational methods that can reconstruct coherent 3D representations from sparse and incomplete ST data.

For medical imaging modalities such as computed tomography (CT) and magnetic resonance imaging (MRI), slice interpolation has long been used to enhance the anatomical information [14–16]. However, direct extension of these approaches to ST is non-trivial. Unlike dense voxel grids with low-dimensional intensity channels, ST data consist of irregularly sampled point clouds associated with high-dimensional gene expression profiles. This combination of spatial sparsity and transcriptomic complexity poses unique computational challenges that are not addressed by conventional interpolation techniques. While several methods have been developed to enhance resolution or impute expression in 2D ST, including both histology-guided and expression-based approaches [17–19], methods specifically designed for reconstructing missing slices and spatial continuity in 3D ST remain limited. Existing 3D interpolation tools often focus on gene expression alone [10, 20, 21], without explicitly modeling spatial structure or cell locations, while recent attempts to infer cell locations are computationally intensive and sensitive to input quality [22]. This gap underscores the need for efficient methods that jointly recover spatial structure and transcriptomic profiles in incomplete 3D ST datasets.

To fill this gap, we present UniST, a unified computational framework for reconstructing dense and continuous 3D ST from sparse serial sections. Rather than introducing new experimental ST technologies, UniST addresses the analytical challenges inherent to current serial section-based datasets. UniST integrates point cloud upsampling for intra-slice densification, optical flow-based slice interpolation for inter-slice densification, and 3D implicit neural representations (INR) for modeling high-dimensional gene expression in 3D space. We evaluate UniST across multiple 3D ST datasets spanning distinct profiling platforms and tissue contexts, demonstrating accurate recovery of missing slices, improved structural continuity, and biologically coherent expression patterns. By alleviating the needs for densely sampled serial sections, UniST provides a generalizable computational solution that complements existing acquisition workflows and supports scalable 3D ST analysis.

## Results

### Overview of UniST

An overview of the UniST workflow is shown in Figure 1. UniST aims to reconstruct a continuous 3D ST land-scape with enhanced resolution from serial 2D tissue sections that are affected by inconsistent point density, tissue loss, and missing slices. To achieve this goal, UniST integrates a generative artificial intelligence (AI) framework composed of three sequential steps. First, UniST applies kernel point convolution with cross-attention layers to perform point cloud upsampling within selected 2D slices, addressing variability in point density across slices. Such heterogeneity commonly arises from cell disruption during tissue processing, technical variability, and associated loss of cell-level information [23]. This step is conceptually analogous to normalization and resolution harmonization procedures widely used in biomedical imaging, where acquisition-related artifacts are corrected prior to downstream analysis. To our knowledge, this represents the first approach to enhance 2D ST resolution while preserving both point density and local geometry. Second, UniST interpolates missing slices by rasterizing individual sections and applying bidirectional optical flow to slice features extracted by a U-Net across multiple hierarchical levels from coarse to fine to estimate intermediate slices. This image-based interpolation is computationally efficient and can perform interpolations on large tissue volumes containing over one million cells within minutes. Finally, UniST performs gene expression imputation by extending the SUICA framework [19] to the 3D setting. A graph autoencoder (GAE) is trained on sparse, high-dimensional gene expression profiles to learn dense, low-dimensional latent representations, and an INR is then fitted to map the 3D spatial coordinates to these representations, which are subsequently decoded to recover gene expression profiles throughout the reconstructed volume.

**Fig 1.**
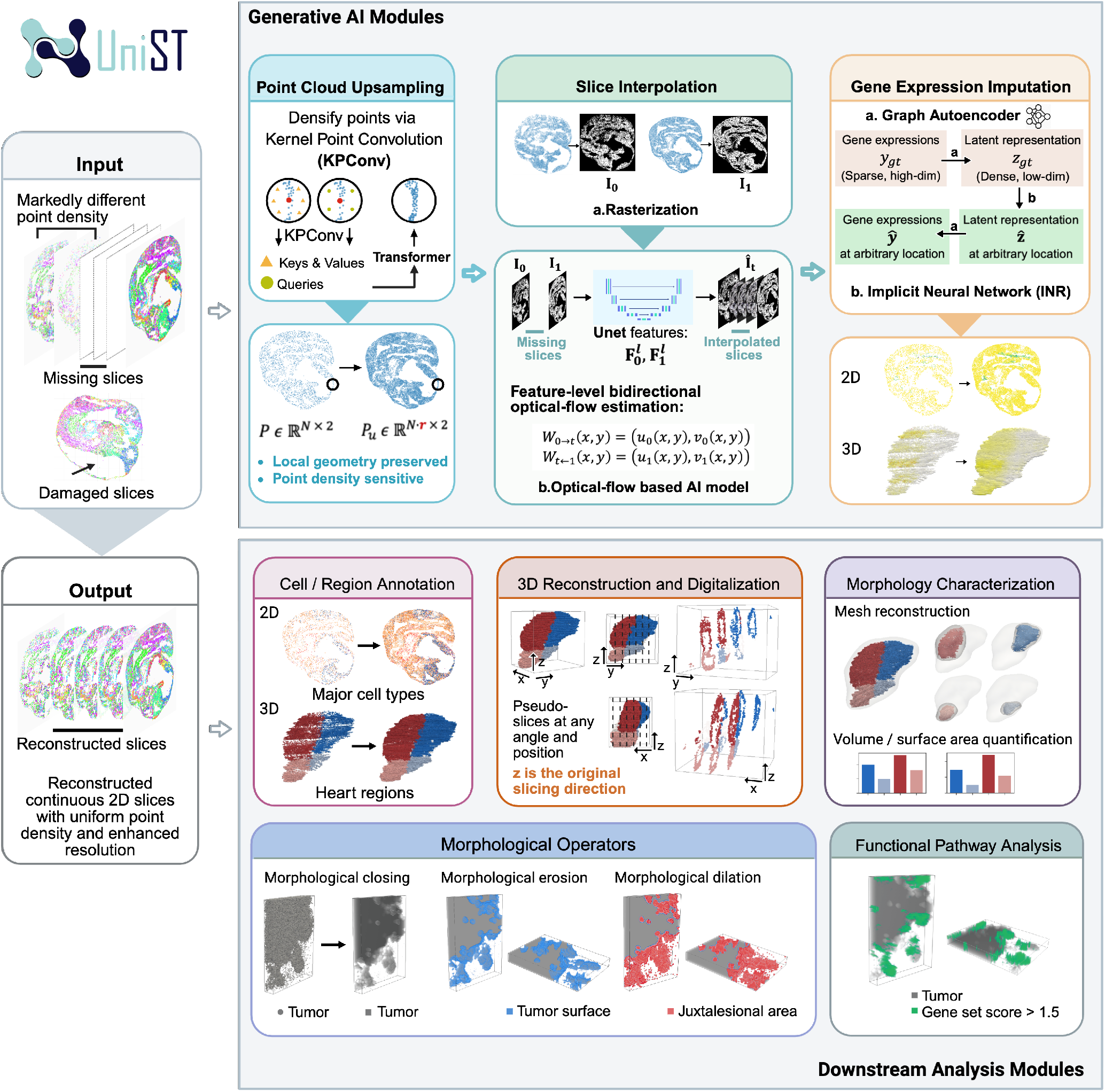
UniST’s pipeline for 3D spatial transcriptomics data reconstruction and downstream analysis. The framework consists of two main modules. The *Generative AI Module* performs point cloud upsampling, slice interpolation, and gene expression imputation to generate continuous and transcriptomically complete 3D ST data. The *Downstream Analysis Module* enables biological interpretation through cell/region annotation, 3D reconstruction and digitalization, morphological operators, quantitative morphology characterization and functional pathway analysis.

After reconstructing continuous 3D ST data with complete gene expression profiles, UniST supports a range of downstream analyses for biological interpretations. These include fast cell- and region-level annotations based on latent representations derived from GAE, pseudo-slice generation along arbitrary planes, and morphological characterization through mesh reconstruction with quantitative assessment of volume and surface area. UniST further provides morphological operators, such as closing, erosion, and dilation, to delineate tumor surfaces and juxtalesional regions, as well as functional analyses to visualize pathway-level spatial patterns.

### UniST evaluation on a 3D mouse embryo dataset

We first applied UniST to a Stereo-seq dataset, MOSTA3D [24], which provides 3D ST maps of a mouse embryo at embryonic day 9.5 (E9.5). This dataset consists of 70 serial sections, each 10 *µm* thick, comprising a total of 646,893 cells with measurements for 17,649 genes. Although 90 sections were generated according to [24], only 70 are available in the public dataset. This dataset exhibits key challenges highlighted in the Introduction, including discontinuities along the z-axis and slice-level heterogeneity (Figure 2a, Supplementary Figure 1), making it a realistic benchmark for evaluating the effectiveness of UniST. Four slices with extensive tissue loss were excluded from this analysis.

**Fig 2.**
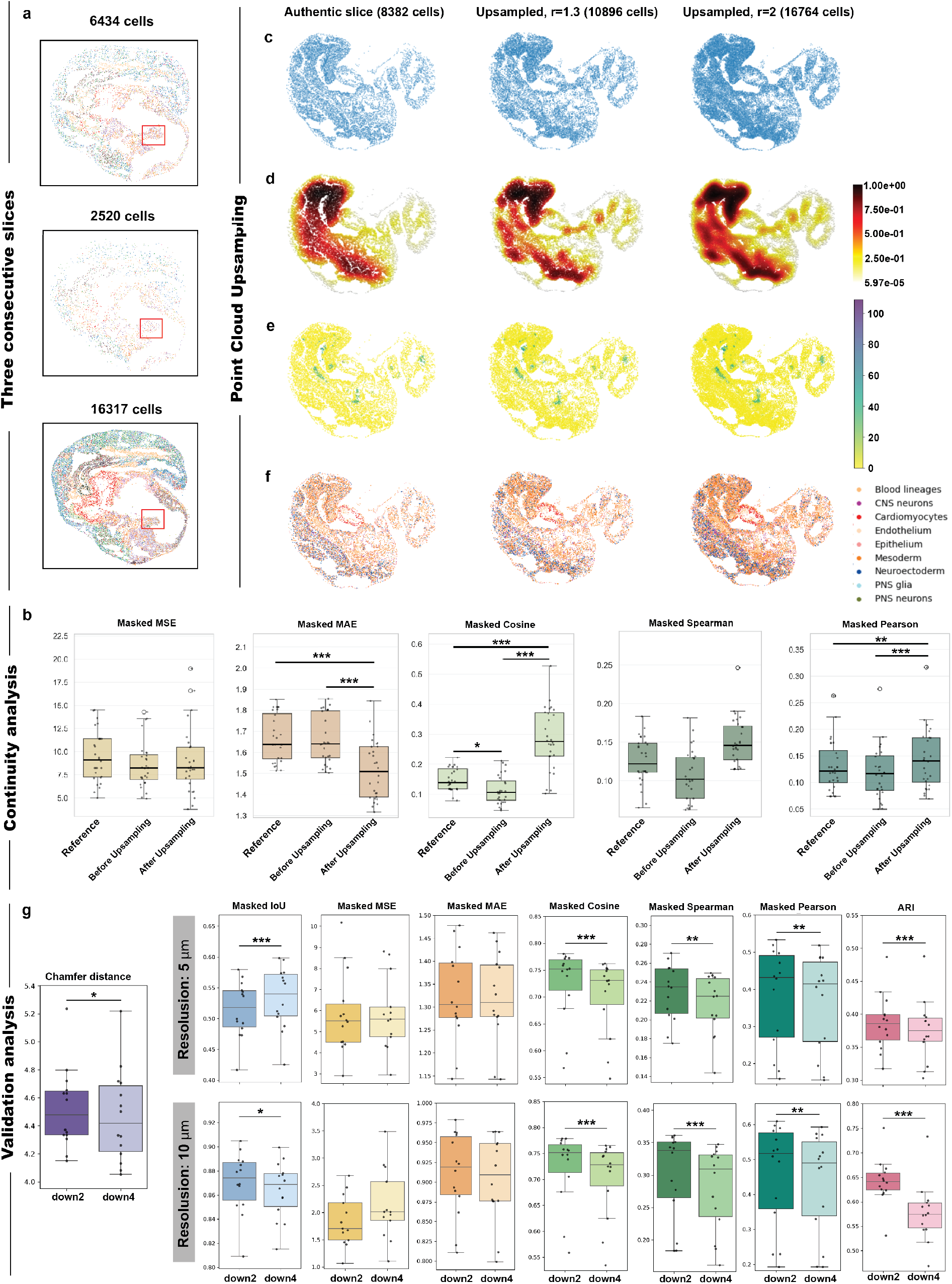
UniST enables shape- and density-preserving point cloud upsampling with experimental validation. (a) Example of three consecutive slices colored by 38 cell types showing heterogeneity across slices (cell type names are shown in Supplementary Fig 1a). (b–e) Comparison of an authentic slice (8,382 cells), the upsampled slice at rate *r* = 1.3 (10,896 cells), and the upsampled slice at rate *r* = 2 (16,764 cells) by UniST. (b) Point cloud visualization. Normalized point density. (d) Gene expression of *Hba-x*, an embryonic α-globin gene expressed in primitive erythrocytes. (e) Cell type annotation of nine major cell types. (f) Masked MSE, MAE, cosine similarity, Spearman, and Pearson correlations computed comparing slices before upsampling and after upsampling with their immediate neighboring authentic slices. Metrics were compared against authentic neighboring slice pairs as references. An illustration of this evaluation strategy is provided in Supplementary Fig 2a. (g) Validation analysis across 14 2D slices. From left to right: structural fidelity, assessed by Chamfer distance (measured by *µm*) and Intersection over Union (IoU) of binned masks (5 *µm* and 10*µm* resolutions); expression fidelity, evaluated by masked MSE and masked MAE; expression correlation, measured by masked cosine similarity, masked Spearman, and masked Pearson correlation; and biological conservation, quantified by ARI. * p < 0.05, ** p < 0.01, *** p < 0.001.

### Point cloud upsampling improves slice-level consistency

To identify slices that are substantially heterogeneous, we quantified the relative differences in spot number and spatial coverage between adjacent sections. Spatial coverage was estimated by binning coordinates at a 5 μm resolution. Slices with a relative difference greater than 1, indicating more than 100% difference from neighboring sections in spot number or spatial coverage, were selected for upsampling and further confirmed by visual inspection. After upsampling with UniST, the relative differences in spot number and spatial coverage between adjacent slices were reduced to below 1 across all sections, indicating improved structural consistency along the z-axis (Supplementary Figure 2a). For slices subjected to upsampling, we further assessed continuity at the gene-expression level by comparing each authentic slice with its preceding and succeeding neighbors and averaging the results (see Supplementary Figure 2b for the schematic illustration). The slice-to-slice continuity was quantified using masked mean squared error (MSE), masked mean absolute error (MAE), masked cosine similarity, masked Spearman’s correlation, and masked Pearson’s correlation, computed across all gene expressions. The mask was defined as a binary matrix derived from the authentic slices, indicating observed spots with expressions (Method). Intuitively, an authentic slice is expected to be more similar to its adjacent slices than the similarity between those two neighbors themselves. However, prior to upsampling, this expectation was violated, i.e., adjacent slices showed higher mutual similarity than comparisons involving the authentic slice (Figure 2b). After UniST upsampling, the similarity between the upsampled slice and its neighbors exceeded that between the two neighbors, demonstrating that UniST improves slice-to-slice continuity along the z-axis.

### Flexible upsampling preserves morphology and expression

Figure 2c-f provide an example slice upsampled at rate *r* = 1.3 and *r* = 2. The use of non-integer factor (*r* = 1.3) highlights UniST’s ability to support arbitrary upsampling rates, rather than being restricted to discrete values. Upsampling at *r* = 1.3 yielded cell counts that are more consistent with adjacent sections. Compared with authentic slices, UniST generated denser point clouds that faithfully preserved both tissue morphology and density distributions (Figure 2c-d). In addition, UniST provided complete gene expression profiles and cell type annotation on the reconstructed point clouds through its gene expression imputation and annotation modules Figure 2e-f.

To quantitatively assess the point cloud upsampling strategy, we performed a systematic evaluation using 14 representative 2D slices spanning diverse anatomical regions. Each slice was randomly downsampled by factors of 2 and 4, after which UniST was applied to recover the original density and gene expression profiles. Reconstructed slices were compared with the authentic slices to assess structural fidelity, expression fidelity, and biological conservation (Figure 2g). Structural fidelity was assessed by Chamfer distance at point cloud level and intersection-over-union (IoU) of binned masks at 5μm and 10μm resolutions. Upsampling from 4X downsampled slices achieved structural fidelity comparable to or even exceeding that from 2X downsampling, except for IoU at 10μm resolution. This trend may be attributable to the model’s pretraining on downsampled inputs at similar spatial scales. [25] (Method). Expression fidelity was evaluated using masked MSE, MAE, cosine similarity, Spearman and Pearson correlations. Biological conservation was quantified by the adjusted Rand index (ARI) based on nine major cell types (see Figure 2f for annotations). As expected, upsampling from 2X downsampled slices generally yielded higher accuracy than from 4X, reflecting the greater information retained in the input data. Evaluations at 10μm resolution were consistently more stable than those at 5μm, indicating increased robustness at coarser spatial scales.

These results demonstrate that UniST enables point cloud upsampling that preserves tissue shape and spatial density while improving slice-to-slice continuity, providing a robust foundation for downstream slice interpolation and 3D reconstruction. To further illustrate the impact of point density on interpolation accuracy, we designed a toy example in which interpolation performance was evaluated under varying density conditions (Supplementary Figure 3; Method). Increased point cloud density consistently improved interpolation accuracy, highlighting the benefit of the upsampling step in UniST.

**Fig 3.**
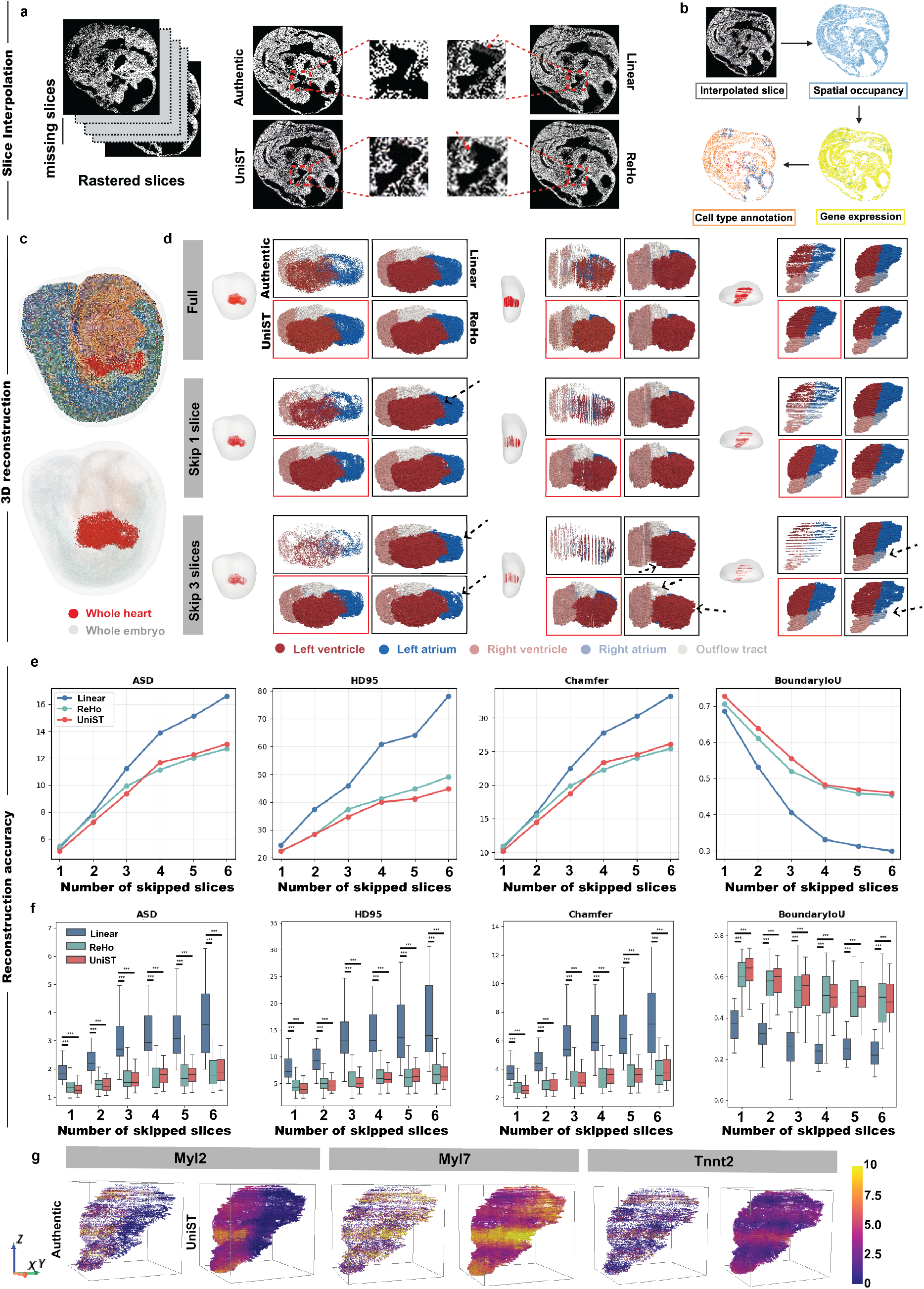
3D heart reconstruction and gene expression imputation with UniST, including comparison with other two interpolation methods and accuracy evaluation when skipping slices. (a) The point cloud was first rasterized into images, and three interpolation methods were compared: linear interpolation, registration-based high-order interpolation (ReHo), and UniST. (b) After interpolation, the rasterized slices were transformed back into occupancy coordinates, followed by gene expression imputation and cell/region annotation. (c) 3D reconstruction of 38 cell types and the whole heart region. (d) 3D reconstruction of five heart sub-regions from the authentic slice stack containing damaged or missing sections, also when skipping 1 slice (10 *µm* interval) and skipping 3 slices (30 *µm* interval) across three interpolation methods. The results are presented from three orthogonal perspectives. (e-f) Quantitatively comparison of the reconstruction accuracy upon the whole heart region at (e) 3D and (f) 2D level across three interpolation methods, measured by ASD, HD95, Chamfer Distance and Boundary IoU. (g) Authentic (left) and imputed (right) 3D gene expression patterns of *Myl2* (ventricular marker), *Myl7* (atrial and early cardiomyocyte marker), and *Tnnt2* (core sarcomeric cardiomyocyte marker) of the whole heart.

### Slice interpolation and whole-tissue reconstruction

Slice interpolation in UniST begins by rasterizing upsampled point clouds into image space, followed by imagebased interpolation. We compared UniST with linear interpolation and a registration-based high-order interpolation method [14] (ReHo; see Method for details). As shown in Figure 3a, linear interpolation introduces ghosting artifacts along tissue boundaries, while ReHo produces smoother but still remains blurred structures. In contrast, UniST provides markedly sharper boundaries and more faithful reconstruction of structural details.

After interpolation, the rasterized slices were transformed back into occupancy coordinates, followed by gene expression imputation and cell/region annotation (Figure 3b). Figure 3 c-d illustrate whole-heart reconstruction with five anatomical subregions. Supplementary Video 1 further provides dynamic visualization of the reconstructed structure. Because only 7 slices were missing within the heart region, visual differences among the three methods were subtle. To further stress-test reconstruction performance and assess robustness to sparse sampling, we manually removed 1 to 6 consecutive slices along the z-axis and reconstructed the missing regions. As shown in (Figure 3d), linear interpolation produced increasingly blocky, low-resolution structures as the number of skipped slices increases. ReHo generates smoother structures than linear interpolation, but occasionally introduces unnatural structural wiggling, potentially due to overfitting (Method). In contrast, UniST maintains smooth and coherent structures without such artifacts.

Quantitatively, we evaluated the reconstruction accuracy of the whole heart region at both the 2D and 3D levels by comparing the authentic slices with their interpolated reconstructions across three interpolation methods. At the 2D level, accuracy was assessed per interpolated slice, whereas at the 3D level, the reconstructed 3D structure was evaluated as a whole. Performance was measured using average surface distance (ASD), 95% Hausdorff distance (HD95), Chamfer distance, and boundary IoU (Figure 3 e-f). These metrics quantify the geometric fidelity and boundary accuracy of reconstructed 2D/3D structures (Method). At the 3D level, linear interpolation consistently underperformed relative to ReHo and UniST across all evaluation metrics. The discrepancy became more pronounced as more slices were skipped. UniST outperformed ReHo in HD95 and boundary IoU, indicating superior preservation of tissue boundary and global structure. For Chamfer distance and ASD, ReHo achieved comparable or slightly better performance when more than three slices were removed.

### Gene expression imputation preserves spatial specificity

Gene expression imputation was subsequently performed on the reconstructed 3D volumes. Figure 3g shows three representative marker genes (*Myl2, Myl7* and *Tnnt2*), illustrating coherent 3D expression patterns along the *z*−axis while preserving localized, gene-specific distributions. Notably, UniST avoids overly smooth predictions and explicitly preserves expression sparsity. For example, *Myl2*, a ventricular cardiomyocyte marker, was predominantly expressed in ventricle. The imputation results by UniST showed negligible expressions in atrium, consistent with known biology. Kernel density estimates (KDE) (Supplementary Figure 4) further show that the highly zero-inflated distribution of *Myl2* expression in authentic data is retained after UniST imputation. In contrast to Guassian process or MLP-based methods, which tend to oversmooth expression patterns and reduce sparsity, UniST preserves non-Gaussian distributions. Both *Myl7* and *Tnnt2* exhibited broad expression across the mouse embryonic heart, with higher expression values observed at the left–right boundary, a spatial pattern consistently recovered by UniST.

**Fig 4.**
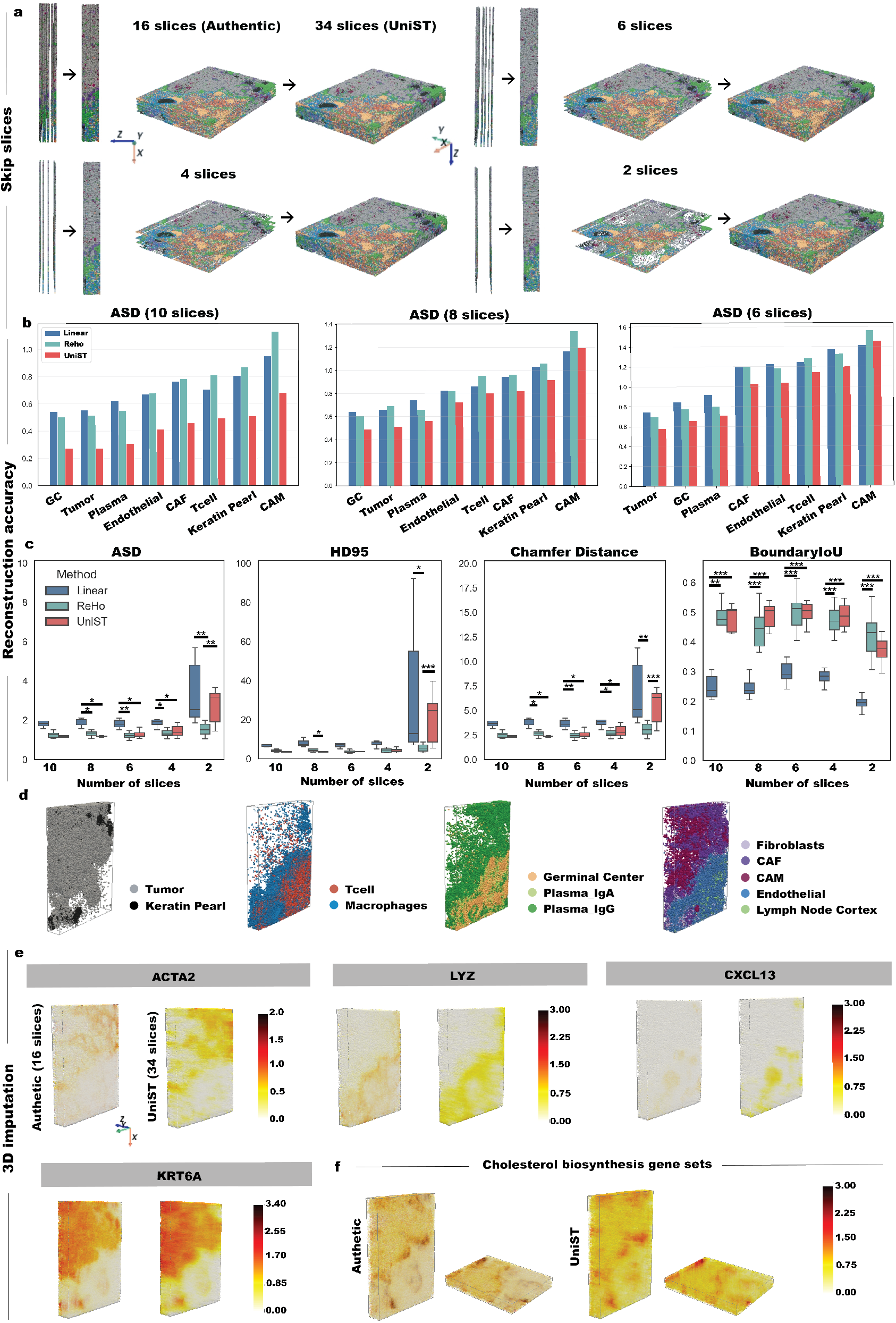
Evaluation of reconstruction accuracy across multiple cellular regions in a human metastatic lymph node, 3D gene expression and functional pathway imputation. (a) 3D reconstructions of authentic data (16 slices) and reconstructed data by UniST (34 slices), as well as representative examples of reconstructions obtained from 6, 4, and 2 slices (after manual skipping). (b) ASD computed at 3D level for reconstruction accuracy using 10, 8, and 6 slices across 8 major tissue regions by three interpolation methods. Regions are ordered from left to right according to increasing ASD values obtained by UniST. (c) ASD, HD95, Chamfer Distance and Boundary IoU computed at 2D level for reconstruction accuracy of the tumor region using 10, 8, 6, 4, 2 slices. (d) 3D reconstruction of major cell-type annotations. (e) Authentic (left) and imputed (right) 3D expression patterns of representative marker genes: *ACTA2* (CAFs marker), *LYZ* (macrophages marker), *CXCL13* (germinal center marker), and *KRT6A* (epithelial marker). (f) Authentic (left) and imputed (right) 3D functional pathway scores of the cholesterol biosynthesis gene set.

### UniST evaluation on a human metastatic lymph node data

Next, we applied UniST on a 3D human metastatic lymph node tissue block by Open-ST [9]. The tissue block measures approximately 4 mm by 3 mm in area and 350 μm in thickness. Similar to Stereo-seq, Open-ST is a sequencing-based ST platform that achieves single-cell resolution. The tissue block was consecutively sectioned at 10 μm thickness per slice. Among them, 19 slices were profiled to obtain ST data for approximately 1.1 million cells with measurements for 28,943 genes. To enable large-area profiling, Open-ST assembles multiple capture areas per slice, which introduces linear cracks in the profiling results (Supplementary Figure 6a). We therefore implemented a semi-automatic crack detection and filling procedure to restore the Open-ST slices (Supplementary Figure 6; Method). In addition, three slices with substantial tissue loss were excluded from the analysis.

### 3D structural reconstruction under sparse sampling

After preprocessing, we systematically evaluated the reconstruction accuracy of UniST in comparison with linear interpolation and ReHo across multiple cellular regions. As in the mouse embryo analysis, we simulated sparse sampling by skipping slices and reconstructed the entire tissue block using 2, 4, 6, 8, 10 slices (out of 34 sections). Figure 4a and Supplementary Figure 7 show the reconstruction results obtained using authentic 16 slices and sparse slice sampling by UniST. Supplementary Video 2 further provides an animation of the 3D reconstruction across major cellular structures. Relatively large and spatially clustered structures, such as tumor (gray) and plasma (green) regions remain recoverable even when only 4 slices were available. To further quantify reconstruction performance across different cellular regions, we computed the same evaluation metrics as before for 8 major cell types. Figure 4b-c and Supplementary Figure 8-9 summarize the 2D and 3D reconstruction accuracy, with tissue regions ordered from left to right by increasing average surface distances. For reconstructions using 6, 8, and 10 slices, UniST consistently achieved higher 3D reconstruction accuracy than both linear interpolation and ReHo across all the tissue regions. When the number of available slices was further reduced to 4 or 2, ReHo outperformed UniST in some regions, a trend that was also observed in the 2D evaluations. This observation is likely attributable to the pre-trained model underlying UniST [26], whose training data did not include such extreme slice sparsity, limiting its ability to extrapolate across very large inter-slice gaps. A similar limitation has been reported in prior work using the same pre-trained model [16]. For this reason, we include ReHo in our package as an alternative, purely algorithmic interpolation option that does not rely on pre-trained models or GPUs.

Across different cellular structures (Figure 4d), germinal center (GC), tumor, and plasma regions consistently exhibited the highest reconstruction accuracy under sparse sampling. As shown in Extended Data Fig1a, these three structures also occupy relatively large fractions of the tissue volume, which likely contributes to their more robust reconstruction. In contrast, smaller or more spatially diffuse structures, such as tumor keratin pearl and cancer-associated macrophages (CAM) showed lower reconstruction accuracy under slice skipping, reflecting their greater sensitivity to missing sections.

### Gene expression and pathway-level imputation in 3D

We next performed 3D gene expression imputation on the reconstructed tissue volumes. Figure 4e shows four representative marker genes, *ACTA2, LYZ, CXCL13* and *KRT6A*, each exhibiting a unique spatial pattern. *ACTA2*, a marker of smooth muscle cells and myofibroblasts, is enriched in stromal regions, including cancer-associated fibroblast (CAF) areas. The imputation results by UniST largely preserved the spatial distribution of *ACTA2* with coherent pattern along z-axis. Similar results were observed for *LYZ* (a macrophage marker), *CXCL13* (a germinal center marker), and *KRT6A* (an epithelial marker). Histograms with KDEs further indicate that the zero-inflated expression distributions are maintained after imputation (Supplementary Figure 10 a-d). In addition to gene expressions, we computed and visualized the functional pathway scores for two representative gene sets, the cholesterol biosynthesis gene set (Figure 4f) and the hypoxia hallmark gene set (Extended Data Fig1b), which were obtained by summing the imputed expression values for genes within each set. Similar to gene expressions, the spatial patterns of the functional pathway scores after imputation closely resemble those observed in the authentic data. Furthermore, histograms and KDEs before and after imputation (Supplementary Figure 10 e-f) indicate that the distributions of the functional pathway scores are more robustly preserved than those of individual gene expression values, likely because the aggregation of multiple genes reduces the sensitivity to gene-level imputation variations [27].

### UniST evaluation on a human gastric cancer data

Finally, we evaluated UniST on a in-house 3D human gastric cancer tissue block generated by our team using Singular G4X, which also provides single cell resolution ST data but uses in situ sequencing. To our knowledge this is the first 3D ST dataset to be reported for this disease context. The tissue block is 4.5 mm × 4.5 mm × 45 μm, and was consecutively sectioned into 9 slices, each 5 μm thick. In total, it provides 3.4 million cells with 335 genes. Slice alignment was conducted using thin-plate spline (TPS) transformation, and initial cell-type annotation was performed based on marker genes. From the data, we observed localized tissue loss occurring at the tissue periphery (Supplementary Figure 11a). Rather than discarding entire sections, we iteratively interpolated missing regions using information from adjacent slices and patched the affected areas, while minimizing perturbation of the original tissue structure (Supplementary Figure 11 b-c). Gene expression imputation and cell-type annotation were then performed by UniST on these patched areas, and the resulting 3D reconstruction is shown in Figure 5a and Supplementary Figure 12, with Supplementary Video 3 providing a dynamic visualization. All the preprocessing details are described in the Method.

**Fig 5.**
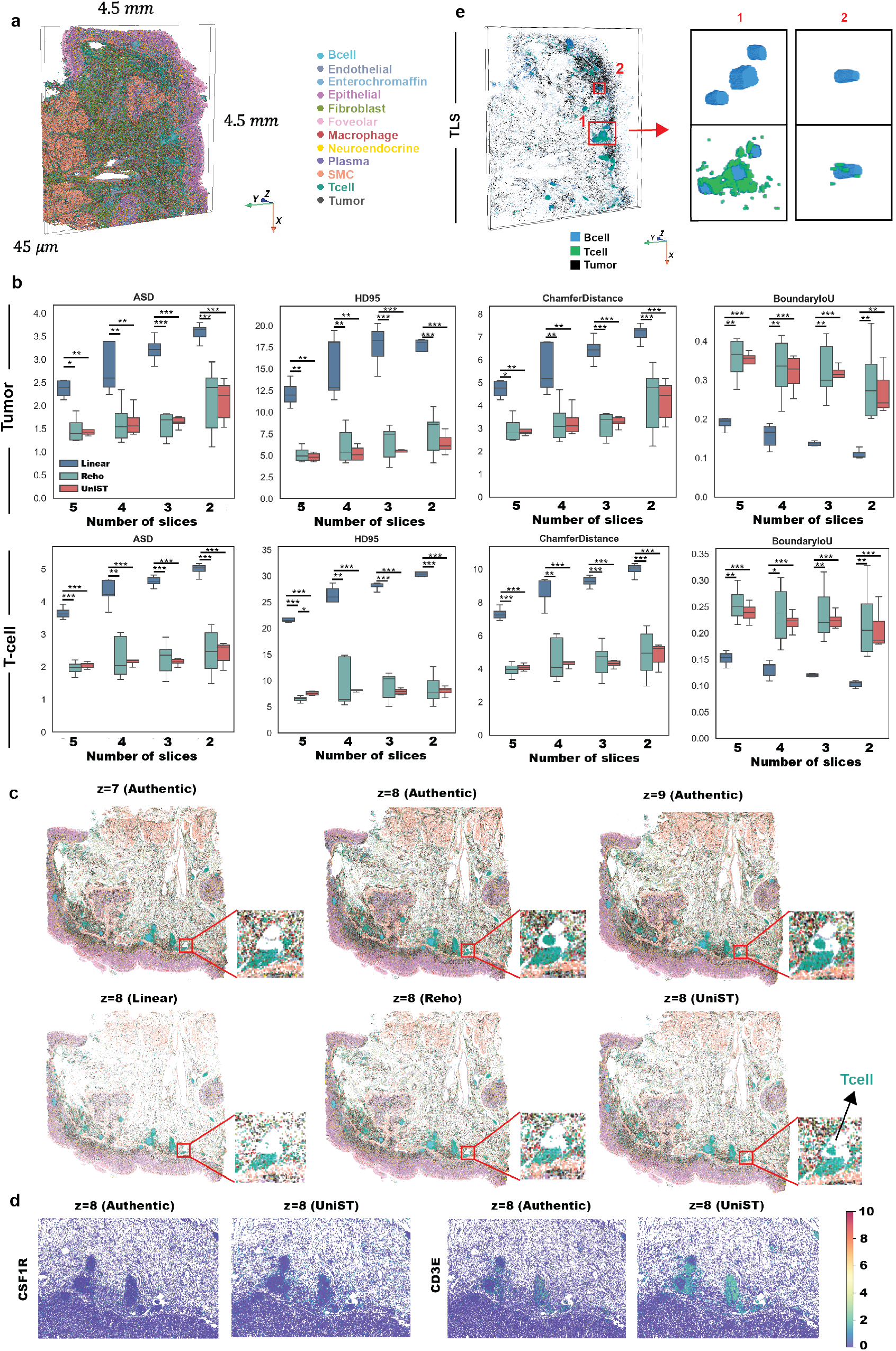
Evaluation of UniST reconstruction accuracy across multiple cellular regions in a human gastric cancer sample and 3D reconstruction of tertiary lymphoid structures. (a) 3D reconstruction of the gastric cancer tissue block colored by annotated cell types. (b) Quantitative evaluation of structural reconstruction accuracy using 2-5 slices, based on ASD, HD95, Chamfer Distance and Boundary IoU computed at 2D level, for tumor (upper) and T-cell (lower) regions. (c) Three consecutive real slices and restored slices by linear interpolation, Reho and UniST. (d) Authentic and UniST-imputed expression patterns of representative marker genes: *CSF1R* (macrophage/monocyte-associated receptor) and *CD3E* (T-cell marker).

In order to quantify the reconstruction accuracy of UniST, we reconstructed tissue volumes using subsets of 2–5 slices without tissue loss and calculated the accuracy metrics across multiple cell type regions. Compared with the previous two datasets, this dataset poses distinct reconstruction challenges, as key cellular structures, including tumor and T-cell regions, are spatially dispersed rather than forming compact clusters. From the results presented by Figure 5b and Supplementary Figure 13, cellular structures with larger spatial extent and occupancy, such as epithelial and smooth muscle cell (SMC) regions, exhibit higher reconstruction accuracy than tumor and T-cell regions with smaller spatial occupancy, which is consistent with previous observations. Across the three interpolation methods, UniST and ReHo consistently outperform linear interpolation. For clustered structure such as SMC, UniST achieved higher reconstruction accuracy than ReHo. But for spatially dispersed structures, UniST exhibits reduced variance rather than higher average reconstruction accuracy across interpolated slices. This observation highlights both the strengths and limitations of UniST, which provides more stable reconstructions of the clustered structures, while remaining sensitive to point cloud density.

We further visualized three consecutive real slices (the 7th, 8th, and 9th slices) and highlighted a small T-cell structure which is absent in the 7th slice but becomes apparent in the 8th slice (Figure 5c). Then the 8th slice was reconstructed using 7th and 9th slices with three interpolation methods. From the results, only UniST successfully recovered this small T-cell structure in the interpolated 8th slice, underscoring its capacity to interpolate structures with gradual spatial transitions along z-axis, which cannot be achieved by algorithmic methods.

Finally, we provide the 3D imputation results of two representative marker genes, *CSF1R* and *CD3E*, visualized on the interpolated 8th slice (Figure 5d). *CSF1R*, a macrophage/monocyte-associated marker, exhibits sparsely distributed expression localized around tertiary lymphoid structures (TLS). UniST preserved these localized signals without excessive smoothing. And similarly, UniST restored the spatial distribution of *CD3E*, which is a T-cell marker. Histograms with KDEs (Supplementary Figure 14) further show that the imputed expression distributions remain zero-inflated, indicating preservation of sparse expression patterns after imputation.

### UniST downstream analysis

UniST facilitates downstream analyses and improves biological interpretations through enhancing the resolution of within and between slices of 3D ST data. The denser reconstructed 3D representation enables pseudo-slice generation at arbitrary orientations, providing greater flexibility of data exploration and visualization. In the mouse embryo dataset, the original data were acquired as serial axial sections, resulting in high resolution in the x–y plane but lower resolution along the *z*−axis. After densification along the *z*−axis, the reconstructed volume was resliced along sagittal and coronal planes (Supplementary Figure 5). The resulting pseudo-slices exhibit clearer structural delineation and improved structural continuity compared with those generated from the original sections, demonstrating the benefit of volumetric densification for multi-planar inspection.

UniST further supports modeling of tissue architecture through voxel-based representation combined with mathematical morphology operators. In particular, morphological closing is applied to rasterized voxel data to connect voxels separated by gaps smaller than a typical cell diameter, yielding continuous tissue structures that more closely resemble those observed in H&E sections. Such morphological operation also bridges the gaps between adjacent 2D sections, improving visualization along the *z*−axis. Extended Data Fig1d presents four representative cellular structures of the human metastatic lymph node after morphological closing, which clearly visualizes the tumor morphology changes along the z axis and the boundary of tumor and T-cell regions. Other major cellular regions after morphological closing are visualized in Supplementary Video 4. Pseudo-slices can be generated at arbitrary angles and positions to inspect the spatial relationships between tumor and T-cell deep inside the tissue block (Extended Data Fig1e). Within the gastric cancer sample, Figure 5e clearly distinguishes between nascent lymphoid aggregates which consist primarily of B cells, and mature TLS architectures, featuring a central B-cell follicle surrounded by a T-cell zone [28, 29]. Beyond structural visualization, UniST allows for 3D rendering of gene expression and functional pathway activity mapped directly onto these structures. For example, the imputed *ACTA2* expression values (>0.75) show strong spatial overlap with CAF structures, while the imputed cholesterol biosynthesis pathway scores (>1.5) are enriched at the tumor boundary (Extended Data Fig1 f-g). To further delineate the invasive margins and juxtalesional areas in the 3D context, UniST applies additional morphological operators, including erosion and dilation, enabling explorations of tumor–microenvironment interactions and proximity-dependent features in the 3D context (Extended Data Fig1h).

In addition to voxel-based analysis, UniST implements mesh representations to model continuous tissue surfaces. Specifically, we extract isosurfaces, which can be viewed as 3D analogues of 2D contours, from the reconstructed voxel data and apply the Marching Cubes algorithm to obtain the mesh reconstructions (Method). Such mesh reconstruction is more accurate when the underlying voxel data are densely sampled, which is achieved by UniST. The morphological features including surface area and volume estimation are then quantified. Supplementary Figure 5 shows mesh reconstructions of the mouse embryo’s heart regions with their surface area and volume measurements. Among these regions, the left ventricle exhibits the largest surface area and volume, whereas the right atrium exhibits the smallest. Collectively, these results highlight UniST’s potential to improve biological interpretation and unlock richer, multi-scale information from 3D ST data.

## Discussion

Transitioning from 2D to 3D spatial molecular profiling is increasingly important for fully capturing the tissue architecture and biological process in their native context [6–8]. At present, most ST datasets are generated through serial sectional strategies [9–12], which are experimentally demanding, costly, and often affected by slice heterogeneity, technical artifacts, and missing sections. Rather than introducing new experimental technologies, UniST addresses these challenges at the computational level by providing a unified generative AI-based framework for reconstructing coherent 3D ST representation from sparse and imperfect serial sections. By enhancing both intra-slice and inter-slice resolution and restoring complete gene expression profiles, UniST enables consistent 3D reconstruction that supports flexible downstream analysis. Across a 3D mouse embryo dataset and two human carcinoma samples, UniST achieved accurate recovery of key anatomical and cellular structures, including the embryonic heart, tumor-immune boundaries, and TLS, under simulated sparse sampling conditions.

Recent methods, including InterpolAI [16] and SpatialZ [22], have demonstrated the potential of interpolation-based strategies for 3D reconstruction, but their performance is largely constrained by the quality and density of the input slices. A key aspect of UniST is the explicit enhancement of in-plane ST resolution prior to slice interpolation. Through the point cloud upsampling module based on kernel point convolution with cross-attention layers, UniST preserves both local geometry structure and transcriptomic density, which are critical for accurate representation of biological context [30]. Extensive evaluation on the mouse embryo dataset shows that this step effectively mitigates slice-to-slice heterogeneity and improves 3D continuity, thereby providing a stronger foundation for subsequent 3D interpolation. In addition, UniST can recover information from regions affected by structural artifacts, such as cracks or localized tissue loss, without discarding entire slices, maximizing the utility of available experimental data.

There are several important improvements and extensions of UniST that can be investigated in future research. First, the current point cloud upsampling and optical flow components rely on pretrained foundation models [25, 26]. Although UniST also provides an algorithmic interpolation method that does not depend on pre-trained models, future domain-specific models trained on large-scale 3D ST datasets-particularly in cancer, where such data remain limitedcould further improve performance. Second, while we demonstrated UniST on sequencing-based and imaging-based ST techniques at single-cell resolution, extending the framework to spot-level ST data still requires additional validation and may benefit from integration with histological images such as H&E data. Beyound transcriptomics, UniST may also be applicable to 3D reconstruction tasks involving other spatial profiling modalities. in particular, for tissue blocks profiled with mixed modalities [13, 31], the framework can inherently handle the resolution discrepancies between different modalities, facilitating the seamless integration of multi-modal datasets into a unified 3D representation. Finally, more downstream analysis can be pursued in the 3D context to further demonstrate the potential applications of UniST, such as cell-cell interaction and RNA velocity.

In conclusion, UniST provides a unified computational framework for reconstructing 3D ST data from sparse serial sections by jointly enhancing intra-slice and inter-slice resolution and supporting versatile downstream analysis. By complementing existing ST acquisition protocols rather than replace them, UniST offers a cost-efficient and generalizable analytical solution for volumetric reconstruction and interpretation of spatial molecular data.

## Method

### UniST Pipeline overview

UniST reconstructs coherent 3D ST structure with complete gene expression profiles and performs downstream analysis for visualization and morphological analysis. The main module of UniST (Generative AI module) consists of three steps: (1) point cloud upsampling to alleviate slice-slice heterogeneity, (2) optical flow–based interpolation to recover the missing slices, (3) INR-based imputation to restore complete transcriptomic profiles.

### Point cloud upsampling

Point cloud upsampling is performed within individual 2D slices to mitigate substantial heterogeneity in point density across serial sections. In the context of ST studies, such heterogeneity arises from variable capture efficiency, tissue damage, and section-specific artifacts, and directly degrades the stability of subsequent slice interpolation and 3D reconstruction. Given a sparse set of 2D points **P** ∈ ℝ^*N ×*2^, the goal is to generate a denser set **P**_*u*_ ∈ ℝ^(*N* ·*r*)*×*2^ while preserving point density and local geometry, where *r* denotes the upsampling rate. Rather than introducing a new upsampling architecture, UniST builds on RepKPU model [25], which was originally developed for geometric point cloud upsampling in computer vision. RepKPU employs Kernel Point Convolution (KPConv) to summarize the local geometric structure of sparse and unordered point sets, serving as an analogue of convolutional neural networks (CNN). While CNN operates on data defined on regular grids (e.g., images), KPConv performs convolutions directly on point clouds by defining a set of learnable, deformable kernel points 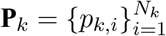, which are uniformly initialized on a circle. Then for each input point *p* ∈ **P**, a circular neighborhood 𝒩 (*p*) is defined by a radius *σ*. This region encompasses a subset of **P** with their corresponding features, denoted as:

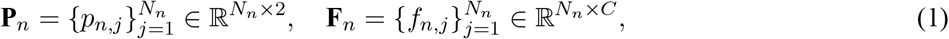

where *N*_*n*_ represents the number of points within 𝒩 (*p*) and *C* denotes the feature dimensionality. For point-only representation, the initial input features simply represent spatial occupancy (*C* = 1). And through subsequent KPConv layers, the feature vectors evolve to high-dimensional latent geometric descriptors, which effectively capture intricate structural information, such as local curvature, density gradients, and manifold topology.

Specifically, KPConv is structured in two stages: spatial correlation and feature transformation. In the first stage, it projects neighboring features **F**_*n*_ onto each kernel point *p*_*k,i*_ via a distance–weighted aggregation:

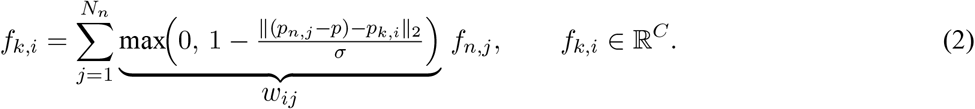

The weight *w*_*ij*_ acts as a spatial correlation coefficient, determining the influence of each neighbor based on its proximity to the *p*_*k,i*_. In the second stage, the projected features *f*_*k,i*_ are multiplied by their corresponding learnable matrices *W*_*k,i*_ ∈ ℝ^*C×C′*^, enabling each kernel point to capture distinct local geometric features. The final output *f*_out_ aggregates these directional insights into a refined latent vector in ℝ^*C′*^ :

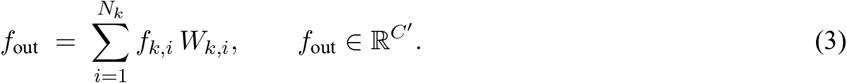

Following feature extraction, RepKPU initializes a set of candidate points within each neighborhood, with their spatial coordinates and latent features as:

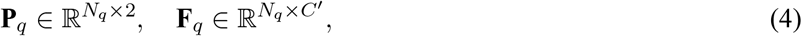

where *N*_*q*_ = *r*, and *r* is the desired upsampling rate. Next, the cross-attention layers are implemented to predict displacement features, which are then mapped through a final regression head to calculate the spatial offset 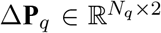. The final upsampled points within the neighborhood are obtained by

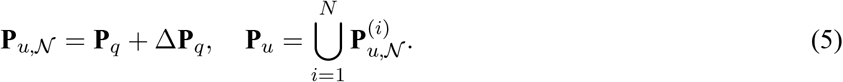

In UniST, we repurposed this geometry-driven upsampling framework as a preprocessing step for biological ST slices, with the specific goal of harmonizing point density across sections prior to 3D interpolation. The original RepKPU model was trained with a fixed upsampling rate *r* = 4 on the PU-GAN dataset, which comprises 120 diverse 3D models utilized to generate 24,000 geometric training patches [25]. To support arbitrary-scale upsampling in our setting, we follow the strategy of first downsampling the point cloud to 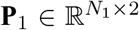, where 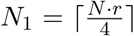 and then applying the pre-trained model to obtain the desired density. The training loss is defined as the weighted sum of Chamfer Distance loss (ℒ_cd_), fitting loss (ℒ_fit_) and repulsive loss (ℒ_rep_), where ℒ_cd_ enforces alignment between upsampled and groundtruth point sets, ℒ_fit_ encourages kernel points to fit the local region and ℒ_rep_ prevents kernel points from collapsing. The detailed mathematical definitions are listed in Supplementary Note 1. Crucially, while the underlying architecture and training objective are inherited from prior work, UniST introduces a new application regime and integration strategy: point cloud upsampling is used here to normalize biological sampling density across heterogeneous ST slices, which in turn improves slice-to-slice consistency and stability of downstream 3D interpolation.

### Slice interpolation

UniST performs inter-slice interpolation using an optical flow-based framework derived from FILM [26], which models apparent morphological “motion” between consecutive sections. FILM was originally developed for video frame synthesis and has recently been adapted to biomedical imaging contexts by InterpolAI [16] for several volumetric imaging modalities. In UniST, we do not introduce a new interpolation architecture; instead, we repurposed this framework for ST data by embedding it within a representation pipeline that bridges sparse point clouds and dense image-like grids. This adaptation, together with the preceding point cloud upsampling and resolution harmonization steps, enables robust interpolation of ST slices while preserving structural integrity and single-cell-scale detail.

Specifically, slice-level point clouds are first rasterized into regular grid representations, which allows the optical flow model to be applied in a controlled and spatially consistent manner. Our experiments show that the combination of density harmonization via point cloud upsampling and careful selection of binning resolution is critical for stable and accurate interpolation; without these steps, direct application of image-based interpolation to sparse ST data leads to structural artifacts and loss of fine-scale detail.

The objective of the interpolation module is to synthesize an intermediate image **Î**_*t*_, between two input images **I**_0_, **I**_1_, where *t* ∈ (0, 1). Unlike traditional interpolation methods that operate in the raw image domain, the framework estimates the ‘motion’ in the latent feature space, which ensures the synthesized slices maintain structural and semantic coherence. To capture both global contexts and local nuances within the tissue sections, it utilizes a U-Net encoder to extract multiscale features at 7 levels from coarse to fine scale from **I**_0_, **I**_1_, denoted as 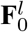 and 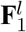. Then the bidirectional optical flow are estimated at each level *l* ∈ [1, 7]. 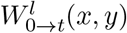 denotes the forward flow field mapping coordinates from 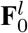 to 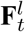:

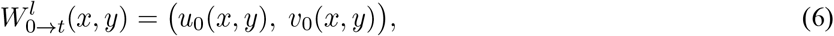

where a pixel located at (*x, y*) in the feature space 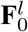 is mapped to the position (*x* + *u, y* + *v*) in the feature space 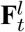. Analogously, the backward flow field 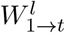 maps feature coordinates from 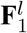 to 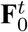 :

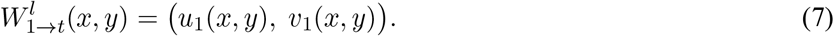

The final synthesis is performed by concatenating features predicted by bi-directional flows and pass them into the U-Net decoder. The model was pre-trained on a largescale dataset consisting of 90k high-resolution videos with large motions ranging up to 120 pixels [26]. The training loss is a weighted sum of pixel-wise *L*1 loss, perceptual loss ℒ_perc_ and style loss ℒ_style_. While the *L*1 enforces structure fidelity, ℒ_perc_ and ℒ_style_ effectively reduce blurriness and enhance edge sharpness. Mathematical definitions are provided in Supplementary Note 1. Within UniST, this interpolation strategy is used to generate an arbitrary number of intermediate slices between any pair of adjacent ST sections. Importantly, the novelty of this component lies not in the optical flow model itself, but in its integration with density-normalized point cloud representations and its role in a unified computational pipeline for reconstructing coherent 3D ST volumes from sparse and heterogeneous serial sections.

### Alternative interpolation method

In addition to the AI-based interpolation module, UniST provides an alternative, purely algorithmic option based on ReHo (registration-based high-order interpolation) [14]. ReHo operates directly in the native coordinate space and estimates bi-directional displacement fields via non-rigid registration, without relying on any pre-trained models. Although the AI-based approach generally achieves superior structural synthesis in most settings, ReHo offers a complementary solution in scenarios with large inter-slice disparities that fall outside the distribution of the pre-trained data, and it eliminates the need for GPU resources.

Given a stack of *P* slices **I**_*k*_ with axial depth *z*_*k*_, ReHo estimates the displacement field 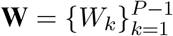 via symmetric, curvature-regularized non-rigid registration. The transformation *p*(*z*, **x**) is then defined as the coordinate mapping that projects a pixel from its original position **x** to a new position at depth *z*. Specifically, *p*(*z*, **x**) = **x** + 𝒮(*z*, **W**), where 𝒮 represents the Hermite spline integration of the optical flow fields across the entire stack. The use of Hermite spline eliminates ‘staircase’ artifacts which are commonly observed from linear interpolation. Then the displacement field **W** is obtained by minimizing the registration energy

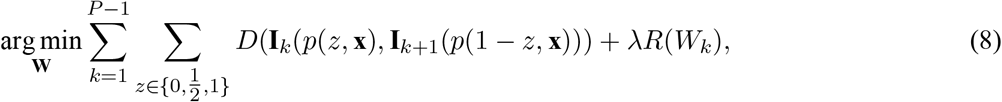

where *D* is the squared *L*_2_ norm and *R* is the curvature regularization which is the second-order derivatives of **W**. The variable 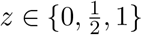 here denotes the normalized axial positions between any two adjacent sections **I**_*k*_ and **I**_*k*+1_. And **I**_*k*_(*p*(*z*, **x**)) means the intensity of **I**_*k*_ at *p*(*z*, **x**). The energy functional is solved using an iterative gradient descent scheme.

After estimating the global trajectories, ReHo synthesizes intermediate slices between **I**_*k*_ and **I**_*k*+1_ by:

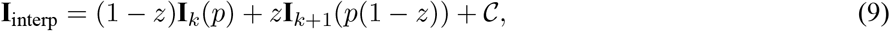

where 𝒞 is the Hermite correction term which integrates global intensity gradients derived from the entire stack to enforce second-order smoothness [14]. Therefore, Hermite interpolation is used twice at both structural and intensity levels.

Within UniST, ReHo serves as a robust, model-free alternative to the learning-based interpolation module, providing a complementary option when computational resources are limited or when the data characteristics fall outside the effective range of pre-trained models.

### Gene expression imputation

Gene expression imputation in UniST is built by extending the SUICA framework [19] from 2D to 3D ST settings. The method is based on INRs models, which use neural networks to map spatial coordinates to their corresponding signals, thereby enabling queries at arbitrary locations within the defined domain. While SUICA demonstrated the effectiveness of this strategy in 2D ST, extending it to 3D introduces additional challenges, including extreme data sparsity, strong anisotropy between lateral and axial resolutions, and pronounced zero inflation in gene expression measurements. UniST addresses these challenges through a set of coordinated architectural and modeling adaptions.

The imputation pipeline first transforms the raw ST measurements **y**_*gt*_ into the latent representations **z**_*gt*_ using a Graph Autoencoder (GAE). This step is motivated by the fact that INRs are better suited to modeling dense, continuous signals, whereas raw ST data are high-dimensional and extremely sparse. The latent representation thus provides a dense and continuous embeddings that could be learned by INR. Specifically, in this step, a connectivity graph is constructed by linking each spot to its k nearest neighbors (kNN) in Euclidean space. A challenge in 3D ST is the discrepancy between lateral (xy) and axial (z) resolutions. To address this, we constructed the connectivity graph using the anisotropic version of kNN. By applying a scaling weight to the z-dimension during distance calculation, we ensured that the model captures more biological information across tissue slices. The loss function to train GAE is masked Mean Square Error (MSE).

The INR is then trained to map 3D spatial coordinates **x** ∈ ℝ^3^ to their corresponding latent representations **z**_*gt*_. To effectively capture high-frequency spatial patterns and fine-scale biological heterogeneity, we utilize a Fourier Feature Network (FFN). In this architecture, input coordinates are projected through a Fourier feature mapping *γ*(**x**) before being processed by a traditional Multi-layer Perceptron (MLP):

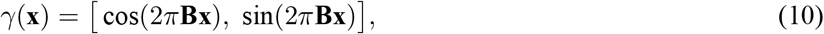

where **B** ∈ ℝ^*m×*3^ is a frequency matrix. To further accommodate the resolution disparity between the lateral and axial dimensions, we implemented an anisotropic frequency sampling strategy for the matrix **B**. The entries of **B** are sampled from a Gaussian distribution, but with distinct variances tailored to each spatial axis:

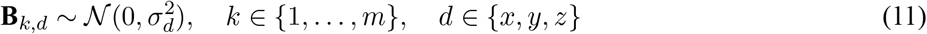

where *σ*_*x*_ = *σ*_*y*_ *> σ*_*z*_. We assigned a set of larger variances to the *x* and *y* directions to generate higher-frequency basis functions, and a set of smaller variances to the *z* direction, yielding smoother, low-frequency bases. This intentional design prevents the model from overfitting the sparse sections. MLPs *f*_*θ*_(·) is then applied on the mapped features:

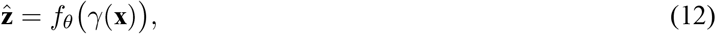

to predict the latent representation. Masked MSE was used as the loss function.

Finally, with the INR parameters fixed, the GAE decoder is fine-tuned to map the predicted latent representations 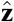 back to gene expression space **y**_*gt*_. The reconstruction loss combines masked MSE, mean absolute error (MAE), and a Dice loss term. The inclusion of the Dice loss is particularly critical given the zero-inflated nature of ST data. Without Dice loss, the model may tend to collapse into only zeros to achieve the lowest reconstruction error. Please see Supplementary Note 1 for detailed mathematical formulations and Supplementary Note 2 for model architectures.

### Toy example to demonstrate the importance of point-cloud upsampling

To illustrate that slice interpolation would be suboptimal without the in-plane resolution enhancement, we designed a toy example using circular structures of diameters 20, 30, and 40 unites. Circular shapes were represented as point clouds, with the 20 and 40 unit circles used for interpolation under both sparse and dense sampling conditions. Three interpolation methods, UniST, ReHo and linear interpolation, were applied, and the reconstructed results were compared with the 30 unit ground truth to evaluate interpolation accuracy. Overall, the results show that denser point clouds lead to more accurate and stable interpolations across all methods (Supplementary Figure 3). ReHo tends to produce slightly smaller circles than the ground truth, while linear interpolation introduced noticeable edge artifacts. In contrast, UniST generates the most faithful reconstructions, and its accuracy improved markedly with increasing point density, demonstrating that the upsampling module effectively enhances in-plane resolution and provides higher-quality inputs for slice interpolation.

### Human gastric cancer data

#### Specimen and ethics

Formalin-fixed paraffin-embedded (FFPE) human gastric tissue from a 50 years-old patient, who developed Hereditary Diffuse Gastric Cancer (HDGC) was obtained through the NIH/NCI biospecimen resources. Sample acquisition and use were conducted under informed consent and approved by the appropriate Institutional Review Board(s), in accordance with institutional policies and applicable international ethical guidelines for biomedical research involving human subjects (NCI protocol #17-C-0043). The specimen was de-identified prior to analysis to protect patient privacy.

#### Serial-section targeted spatial multiomics on the Singular G4X platform

To enable 3D reconstruction, a representative area of interest of 10 × 10 mm containing tumor cells, their microenvironment and at-risk gastric mucosa (non-tumor mucosa), was selected with pathologist guidance. Then, nine consecutive 5-µm-thickness sections from the same FFPE tissue block were collected as serial sections, and transferred for a single run in a two-column layout slide (5 sections in one column and 4 sections in the other column, with one gastric adenocarcinoma as control) and profiled using the Singular Genomics Systems G4X Spatial Sequencer with a targeted in situ spatial transcriptomics workflow (custom 360-gene panel). In parallel, a 17-plex protein panel was applied to enable spatial protein readout as part of the targeted multiomic assay. The G4X workflow supports serial-section, multi-section layouts for 3D-oriented reconstruction, with tissue regions processed in standardized run configurations. Tissue sections were transferred using the vendor’s workflow. Targeted transcript detection relies on padlock probe recognition followed by rolling circle amplification (RCA) to generate localized signal within tissue. Imaging and in situ sequencing were performed on-instrument. In addition to molecular readouts, the platform provides an integrated morphology channel via fluorescent H&E (fH&E) for structural context and quality assessment.

#### Primary processing, single-cell matrix generation, QC, clustering, and annotation

Primary processing was performed using the G4X analysis suite to generate per-cell outputs, including transcript/protein feature quantification, cell segmentation, spatial coordinates, and run-level QC summaries. Exported single-cell outputs (count matrices and spatial metadata) were used for downstream analyses in Seurat (v5.2.0).

Cells with fewer than 10 detected genes were excluded. Additional low-quality cells were removed based on the absence of canonical lineage marker expression and/or other dataset-specific QC criteria. Normalization and variance stabilization were performed using SCTransform, retaining all genes for downstream modeling. Principal component analysis (PCA) was performed using 30 components, followed by construction of a k-nearest-neighbor (kNN) graph and Louvain clustering (resolution = 0.5). Cluster marker genes were identified using FindAllMarkers, and cell types/states were assigned based on canonical lineage markers and established gastric epithelial, immune, and stromal signatures. Where appropriate, assigned labels were mapped back to spatial coordinates for morphology-consistent validation across serial sections.

#### Spatial registration of serial sections using fH&E landmarks and thin-plate spline warping

To integrate cell-level spatial coordinates across serial sections for 3D reconstruction, we performed section-to-section spatial registration using the fluorescent H&E (fH&E) morphology images produced by the G4X workflow. Because the fH&E images and the per-cell spatial coordinates share the same coordinate frame, landmarks selected on fH&E directly correspond to the cellular coordinate system. For each pair of adjacent sections, we manually selected 20 corresponding anatomical landmarks on the fH&E images (e.g., distinctive gland boundaries, stromal–epithelial interfaces, vessel edges, and other reproducible microanatomical features) to define point correspondences between the moving and fixed sections. Using these matched landmark coordinates, we fit a 2D thin-plate spline (TPS) transformation that minimizes non-rigid bending while matching the selected landmark pairs. TPS fitting and coordinate transformation were implemented in Python using RBFInterpolator (SciPy v1.17.0). The fitted TPS warp was then applied to all percell spatial coordinates (cell centroids) from the moving section to map them into the fixed section coordinate space. To register all sections into a common reference frame, we designated the bottom (reference) section as the target coordinate system and performed registrations sequentially in a layer-by-layer manner: each section was registered to its immediate neighbor using TPS, and the resulting transforms were composed to map cells from each section into the bottom reference section. Registration quality was assessed by visual inspection of fH&E overlays and by evaluating residual landmark alignment errors after warping.

### Slice preprocessing

We benchmarked UniST on three real 3D datasets spanning distinct profiling techniques, biological contexts, and spatial complexities. Each dataset exhibits a different challenge, representing common data issues encountered in 3D ST: the 3D mouse embryo dataset has pronounced slice-to-slice heterogeneity, the human metastatic lymph node dataset contains linear cracks, and the human gastric cancer dataset shows localized tissue loss. We addressed slice-level heterogeneity by point cloud upsampling module and applied dedicated preprocessing steps to linear cracks and localized tissue loss, as described below.

#### Filling linear cracks

To enable spatial profiling of large tissue areas, Open-ST [9] assembles multiple capture areas to profile a single tissue section, which introduces linear cracks on the data. We developed a semi-automated pipeline to address this stitching-induced discontinuities (Supplementary Figure 6 a-c). To facilitate automated crack detection, we first rasterized the ST data into an occupancy grid and applied an initial rotation to minimize the angular deviation (*<* 30^°^) between the linear cracks and the *x*-axis manually. We then performed a global row-wise search to detect cracks, where rows with occupancy levels significantly lower than those of their local neighborhood were flagged as candidate crack regions. Next, we extracted local bands surrounding the candidate rows and conducted a column-wise refinement by evaluating occupancy continuity perpendicular to the crack axis, enabling precise delineation of the slope of each linear crack. The identified linear gaps were filled using a Kernel Density Estimation (KDE)-based sampling strategy. By estimating the 2D probability density function in the proximal neighborhood, we generated synthetic points to restore the local point density. As shown in Supplementary Figure 6 d-e, this pipeline produces a continuous point cloud with coherent gene expression values, cell-type regions and tumor–immune boundaries, and provides a foundation for downstream slice interpolation.

#### Restoring local tissue loss

The gastric cancer dataset was generated by the Singular G4X platform, with each tissue section being 5 *µm* thick. Due to the physical fragility of these thin sections, we observed localized tissue loss at the tissue periphery (Supplementary Fig. 11a). Rather than discarding entire sections, we iteratively interpolated missing regions using information from adjacent slices and patched the affected areas while minimizing perturbations to the original tissue structure (Supplementary Fig. 11 b-c). Specifically, we first computed the density difference map between the original slice and the interpolated slice constructed from adjacent ones. Candidate missing regions were identified by thresholding the density difference. To avoid modifying well-profiled regions, we defined a protection zone based on the tissue boundary and removed patch candidates within this zone. Finally, we filled the original slice by the remaining patch to restore the 2D section.

### Computational efficiency

UniST leverages pre-trained models for point cloud upsampling and slice interpolation. When deployed on an NVIDIA L4 GPU via Google Colab, the runtime for point cloud upsampling and slice interpolation is under 60 seconds. The gene expression imputation was performed on a standard 6-core CPU workstation. Its runtime varies across datasets: approximately 3.5 hours for the mouse embryo heart, 10.5 hours for the human gastric cancer sample, and 13.5 hours for the metastatic lymph node. In all cases, the GAE training phase accounts for the majority of the execution time.

### Downstream analysis

#### Cell type annotation

For cell type annotation, UniST provides a lightweight strategy by transferring the annotations on original data points to synthetic coordinates through a joint *k*-NN search across both transcriptomic and spatial domains. Specifically, for each newly generated point *u*, we identify its *k* nearest neighbors within the original labeled points by computing a joint distance metric that balances latent gene expression embeddings (derived from the GAE) and spatial coordinates.

Let **p**_*i*_ ∈ ℝ^*d*^ (*d* ∈ {2, 3}) denote the coordinates of *N* original points, and 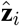 to be their corresponding latent embeddings. Each original point is associated with a cell type label 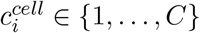. For a synthetic point *u* at coordinate **p**_*u*_ with embedding being 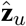, the distance metric to an original point *i* is defined as:

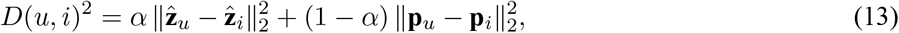

where *α* ∈ [0, 1] modulates the contribution of expression similarity versus spatial proximity. The optimal choice of *α* depends on the characteristics of the dataset. We used *α* = 0.1 for all three datasets in this study. Final cell type assignment for point *u* is determined by majority voting across the labels of its *k* nearest neighbors *N*_*k*_(*u*) based on *D*(*u, i*).

#### Morphological operations for tissue architecture modelling

We employed 3D mathematical morphology operators on the voxelized representations of the ST data. Let 𝒱 ⊂ ℤ ^3^ represent the binary voxel set of a reconstructed tissue component, and *B* denote a spherical kernel with radius *r*. We used *r* ≈ 5 *µm* to approximate the average radius of a single cell. The dilation of 𝒱 by *B*, denoted as 𝒱 ⊕ *B*, expands the tissue boundaries by one cell-width in every direction. The erosion, denoted as 𝒱 ⊖ *B*, shrinks the tissue boundaries by one cell-width. The morphological closing, defined as

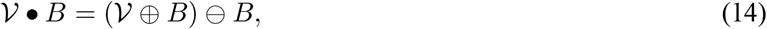

performs dilation and subsequent erosion to bridge voxel gaps and hollow spaces smaller than a cell diameter, yielding a coherent manifold for mesh reconstruction. The 3D boundary, representing the tissue’s leading edge, is delineated by computing the set difference between the original tissue volume and its erosion:

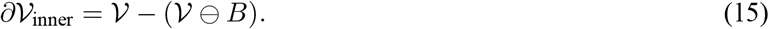

The outer zone, representing the space directly adjacent to the tissue surface, is obtained via:

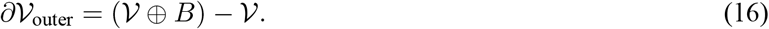

Through the recursive application of these operators or by varying the radius of the kernel *B*, distinct, nested layers of the tumor microenvironment can be defined, facilitating the systematic profiling of molecular gradients across the tumor–stroma transition.

#### Pseudo-slice generation and 3D visualization

For pseudo-slices generation, we developed a computational framework for generating multi-planar virtual sections at arbitrary orientations. Mathematically, given a reconstructed 3D point cloud *P* and a user-defined normal vector **n**, a stack of *N* slices is generated by defining a sequence of plane origins 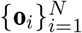 along the axis of **n**. The sampling range is determined by projecting the point cloud onto the normal vector to identify the volumetric extrema (*t*_*min*_, *t*_*max*_):

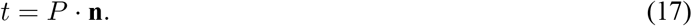

A boundary margin is then applied such that the slice origins are constrained within [*t*_*min*_ + margin, *t*_*max*_ − margin]. Each virtual slice is extracted by querying all points within a specified distance *w* (slice width) from the plane:

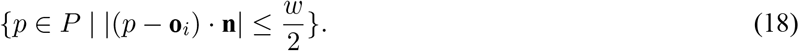

Mesh reconstruction based on Marching Cubes [32] with the quantitative measurements of surface area and volume were conducted based on the code of Spateo [10]. The point cloud data in this study was visualized by PyVista [33]. The voxel data in this study were visualized by ParaView [34].

### Evaluation metrics

To evaluate the expression fidelity and continuity, we employed masked MSE, MAE, cosine similarity, Spearman and Pearson correlation, with the mask defined on the voxelized representation of the authentic slices, indicating observed spot–gene pairs. Mathematically, the masked MSE is defined as:

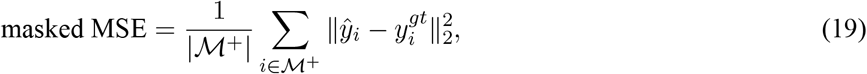

where ℳ^+^ represents the set of indices for observed spots, |ℳ^+^| denotes the total number of these observed spots, and 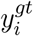 is the observed expression vector for spot *i*. Masked MAE is defined as:

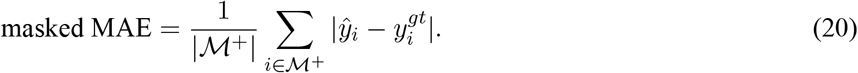

Masked Cosine Similarity is defined as:

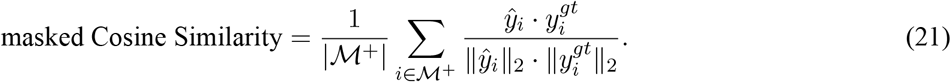

For masked pearson and spearman correlation, spots with fewer than two observed genes or zero variance in expression were excluded from the calculation. Let |ℳ^+∗^| denote this filtered subset of spots. The masked pearson correlation is defined as:

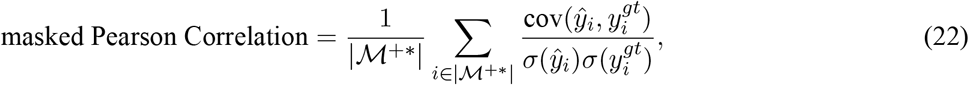

and the masked spearman correlation is:

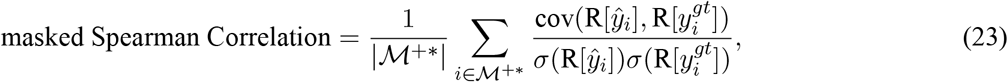

where R[·] denotes the rank operator.

To quantitatively assess the reconstruction accuracy, we employed four geometric metrics in both 2D and 3D space: average surface distance (ASD), 95% Hausdorff distance (HD95), Chamfer distance, and boundary intersection-over-union (IoU). Given two point sets *A* (prediction) and *B* (ground truth), ASD is defined as

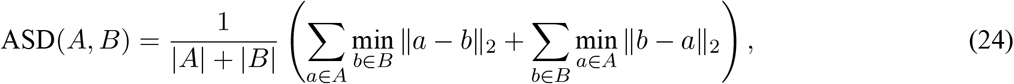

which measures the mean geometric deviation between structures, reflecting overall shape accuracy. HD95 is defined as

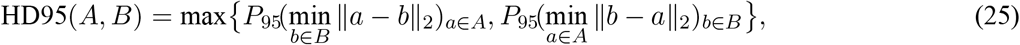

where *P*_95_(·) denotes the 95th percentile operator. It thereby quantifies the worst-case structure deviation while remaining robust to outliers. Chamfer distance is defined as

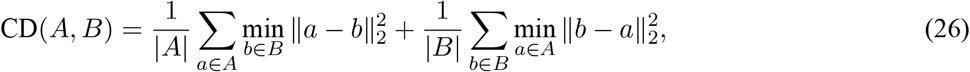

which evaluates the bidirectional point-wise similarity, capturing dense geometric correspondence. Finally, the boundary IoU is defined as

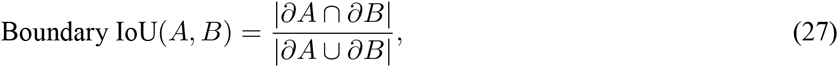

where *∂A* and *∂B* denote the boundary obtained via morphological operators. Boundary IoU focuses on edge alignment, offering a more sensitive measure of contour preservation than standard IoU.

## Supporting information

All in one

## Data availability

The 3D mouse embryo data is available at https://spateodata.aristoteleo.com/. The 3D human metastatic lymph node data is available at GEO (accession numbers GSE251926). The 3D human gastric cancer data will be made available after publication acceptance.

## Code availability

**UniST** is released as an open-source Python library on GitHub at https://github.com/lanshui98/UniST. Tutorials of **UniST** with implementation details are available at https://unist-tutorial.readthedocs.io/en/latest/.

## Acknowledgments

LS, YY, LL, and ZL were partially supported by the Coordination and Data Management Center of the Translational and Basic Science Research in Early Lesions Program (U24CA274212). ZL and LS were also supported by the National Institute of General Medical Sciences (R35GM159819). YL, YD, and LW were partially supported by the National Cancer Institute (U01CA294518, U01CA264583, R01CA266280) and Break Through Cancer. LW also acknowledges research support from the James P. Allison Institute and the Institute for Data Science in Oncology at The University of Texas MD Anderson Cancer Center.

## Author contributions

ZL and LL conceived the project with inputs from LS and LW. LS implemented the methods, analyzed data and prepared the manuscript. YL, LJIC, JRC, YD, WL, KK, RB, JM, REW, KC, THH, PFM, JD, LMSS, LW generated the human gastric cancer 3D ST dataset. YL preprocessed the human gastric cancer 3D ST dataset. All authors reviewed the draft and approved the current version.

**Extended Data Figure 1.**
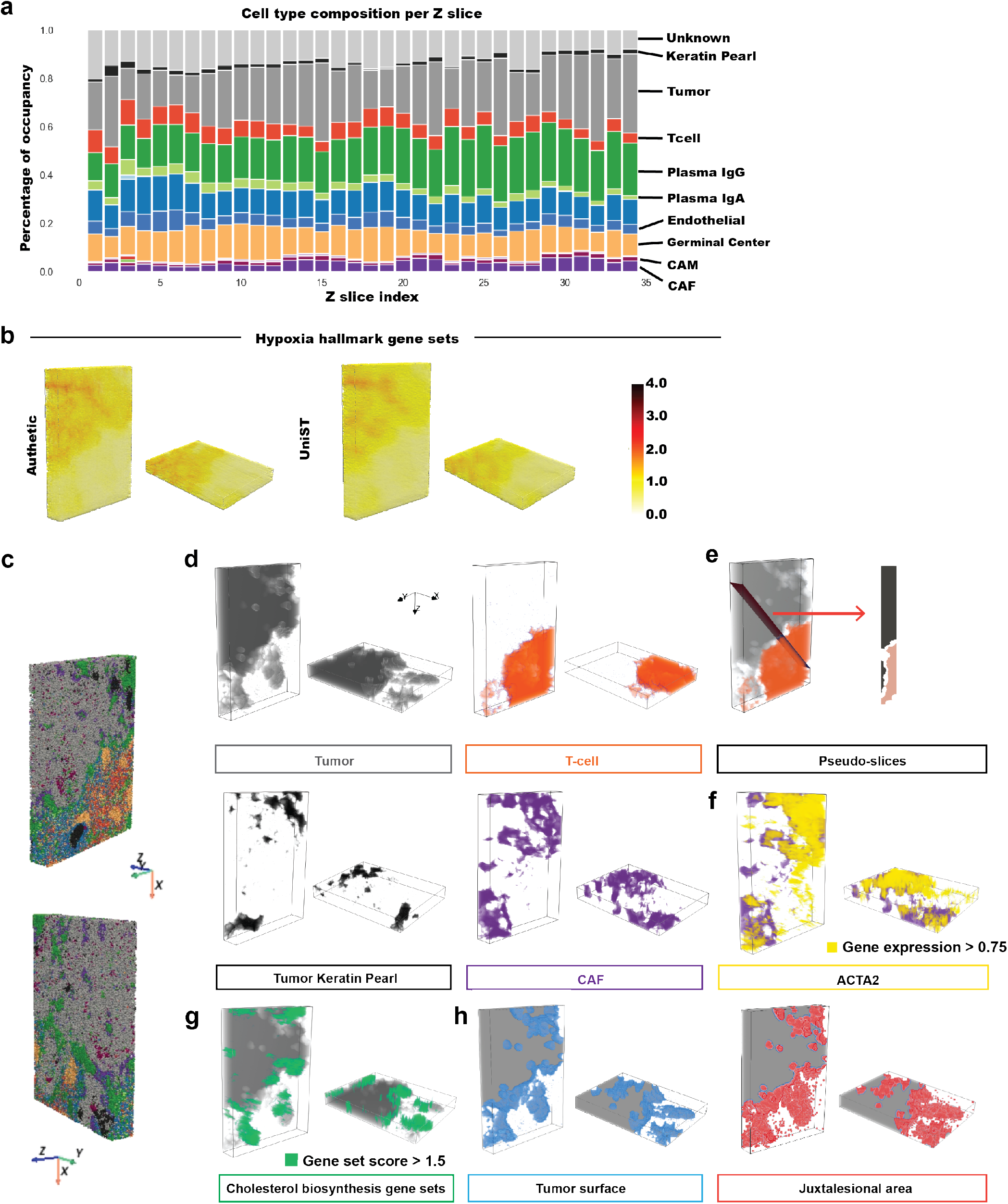
Slice-wise composition, pathway-level imputation and downstream visualizations of the 3D human metastatic lymph node. (a) Cell-type composition of major cell types across reconstructed 34 slices by UniST, illustrating slice-to-slice continuity. (b) Authentic (left) and imputed (right) 3D functional pathway scores of the hypoxia hallmark gene set. (c) 3D reconstruction of major cell types. (d) 3D rendering of tumor, T-cell, tumor keratin pearl and CAF after morphological closing. (e) Pseudo-slice generation of the spatial relationship between tumor and T-cells. (f) 3D expression pattern of ACTA2 visualized with CAF, presented by the isosurface of expression values greater than 0.75. (g) 3D distribution of the cholesterol biosynthesis gene set score visualized with tumor, presented by the isosurface of score values greater than 1.5. (h) 3D rendering of tumor surface based on morphological erosion (left) and juxtatumoral regions based on morphological dilation (right).

