## Supplementary material for "UniST: A Unified Computational Framework for 3D Spatial Transcriptomics Reconstruction": All in one

### Supplementary Information

3.4 Supplementary Figure 14. Gene expression distribution before and after UniST imputation. 16

**Supplementary Notes** **17**

**4 Supplementary Note 1. Training loss functions.** **17**

**5 Supplementary Note 2. Model architectures and hyperparameters for gene expression imputation.** **19**

### Supplementary Figures

### 1 List of Figures for 3D mouse embryo data by Stereo-seq

#### 1.1 Supplementary Figure 1. Characterization and evaluation of slice heterogeneity.

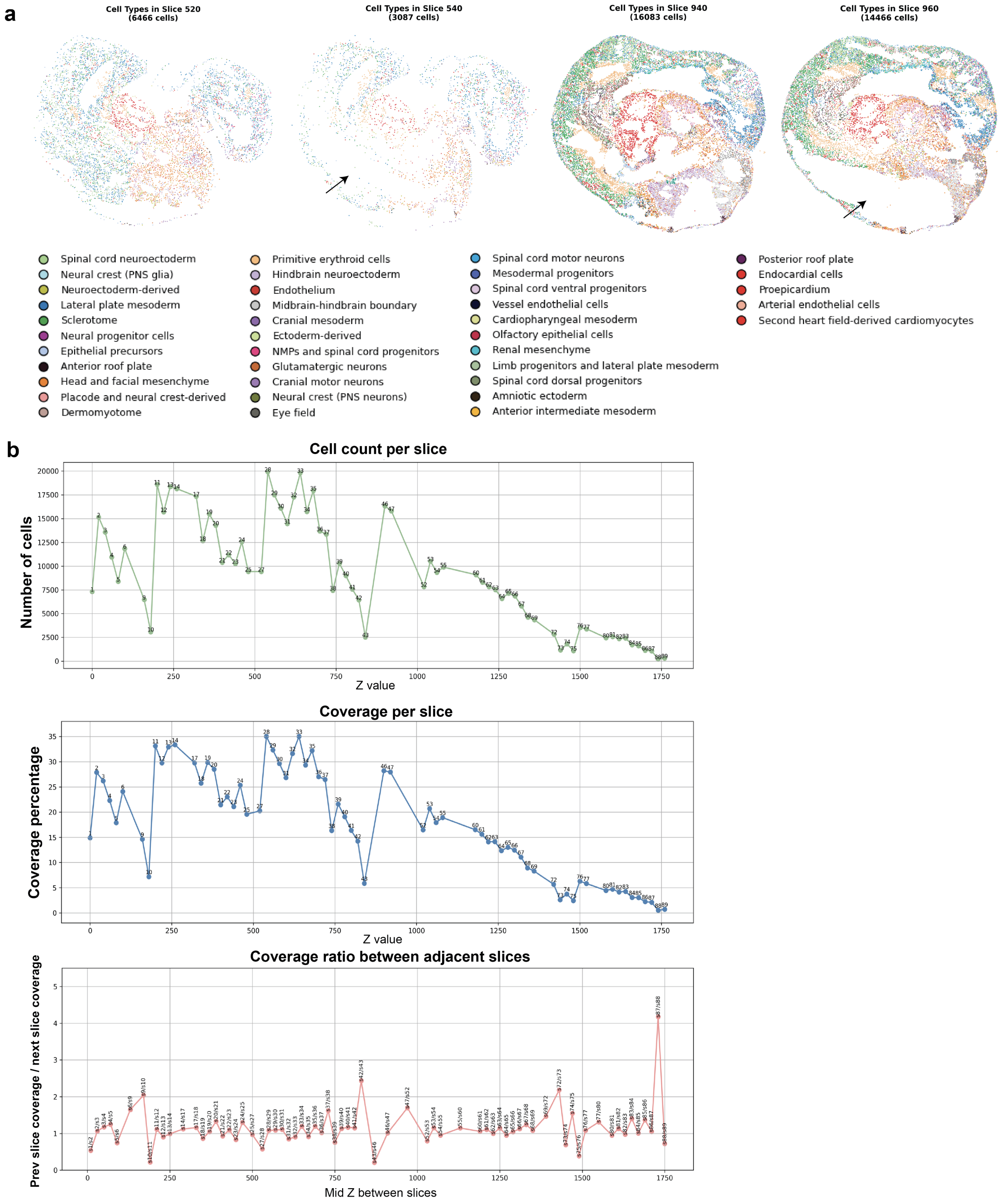

(a) Example of two slices with tissue loss (with arrow) with their adjacent slices (without arrow) colored by 38 cell types. (b) Heterogeneity across 2D slices. From top to bottom: cell count per slice, coverage per slice after binning at 5 $\mu m$, and coverage ratio between adjacent slices.

#### 1.2 Supplementary Figure 2. Slice continuity evaluations.

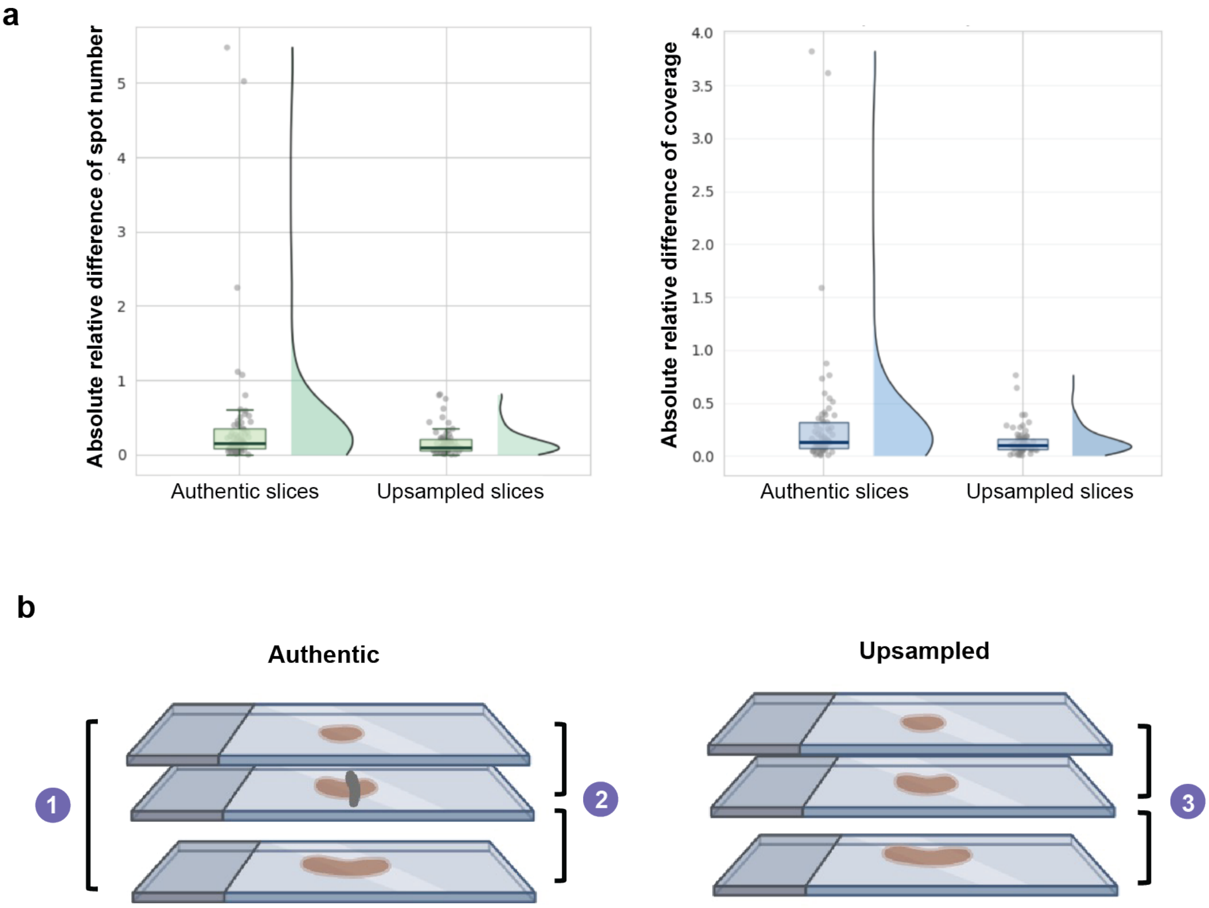

(a) Slice-to-slice continuity quantified by the absolute relative difference in spot number and occupancy coverage at 5  m resolution between adjacent slices before and after upsampling across all sections. (b) Illustration of slice continuity evaluation strategy for Fig 2g. 1. Compare authentic neighboring slice pairs as the reference. 2. Compare authentic heterogeneous slice with its neighboring slices and take average. 3. Compare upsampled slice with its neighboring slices and take average.

#### 1.3 Supplementary Figure 3. Simulation of circular point-cloud structures for benchmarking interpolation accuracy.

a

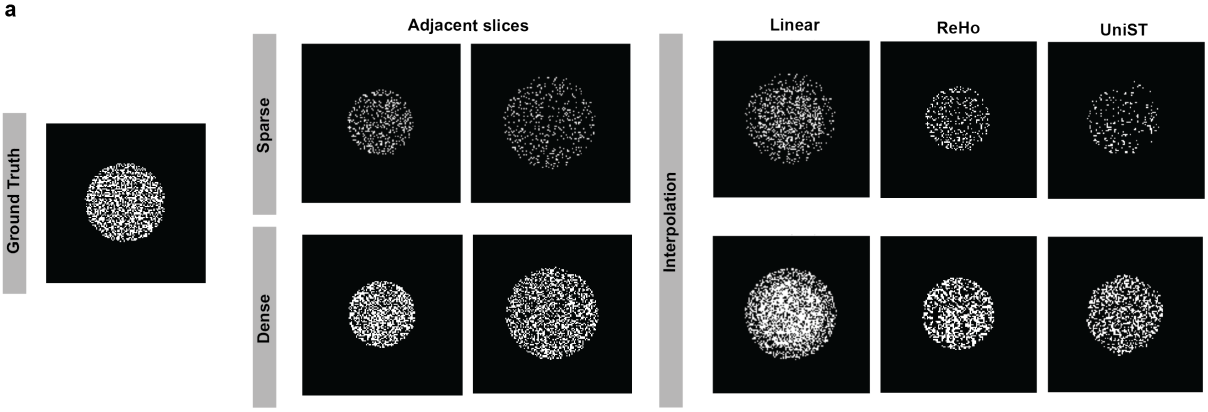

b

| Method | ASD(↓) | HD95(↓) | Chamfer(↓) | Boundary IoU(↑) |
|---|---|---|---|---|
| **Sparse** | | | | |
| Linear | <u>0.97</u> | **4.12** | <u>1.94</u> | **0.79** |
| UniST | **0.78** | <u>4.47</u> | **1.58** | <u>0.78</u> |
| ReHo | 1.24 | 5.39 | 2.49 | 0.67 |
| **Dense** | | | | |
| Linear | 0.86 | 5.00 | 1.73 | 0.75 |
| UniST | **0.67** | **2.00** | **1.35** | **0.93** |
| ReHo | <u>0.71</u> | <u>2.24</u> | <u>1.42</u> | <u>0.87</u> |

(a) Simulation of circular structures with diameters of 20, 30, and 40 units represented as point clouds. Circles with diameters of 20 and 40 were used for interpolation under both sparse and dense sampling conditions. Three interpolation methods were applied, and the reconstructed results were compared with the ground truth to evaluate interpolation accuracy. (b) The numeral results of (a). **Bold** figures are best scores and <u>underlined</u> figures are second-best.

#### 1.4 Supplementary Figure 4. Gene expression distribution before and after UniST imputation.

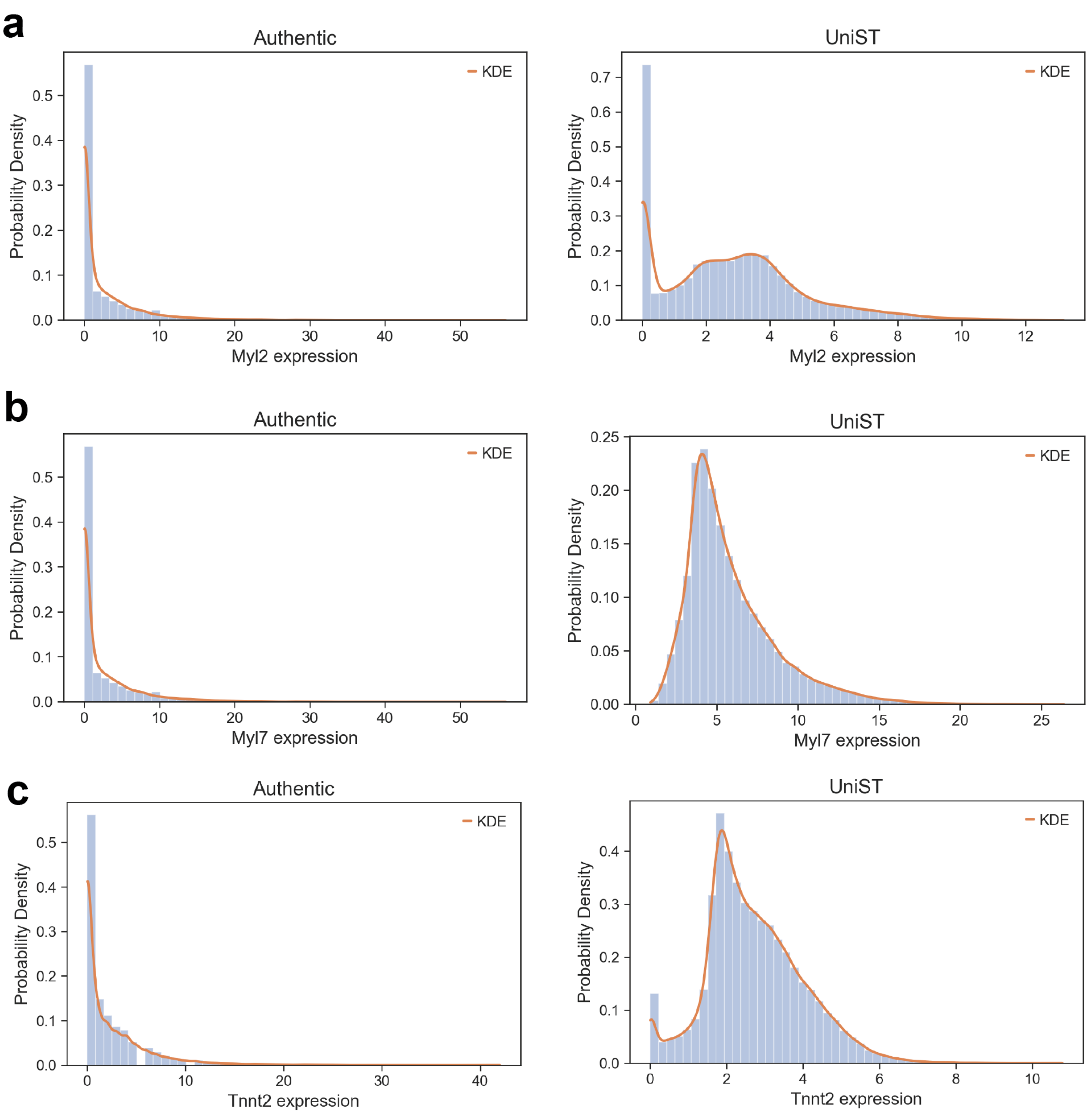

Histograms with kernel density estimates (KDE) for the distribution of expression values for Myl2 (a), Myl7 (b), and Tnnt2 (c) in the authentic data (left) and after UniST-based imputation (right). Histograms were computed using 50 equally spaced bins.

#### 1.5 Supplementary Figure 5. 3D mesh reconstruction of heart regions, morphological quantification and pseudo-slice generation.

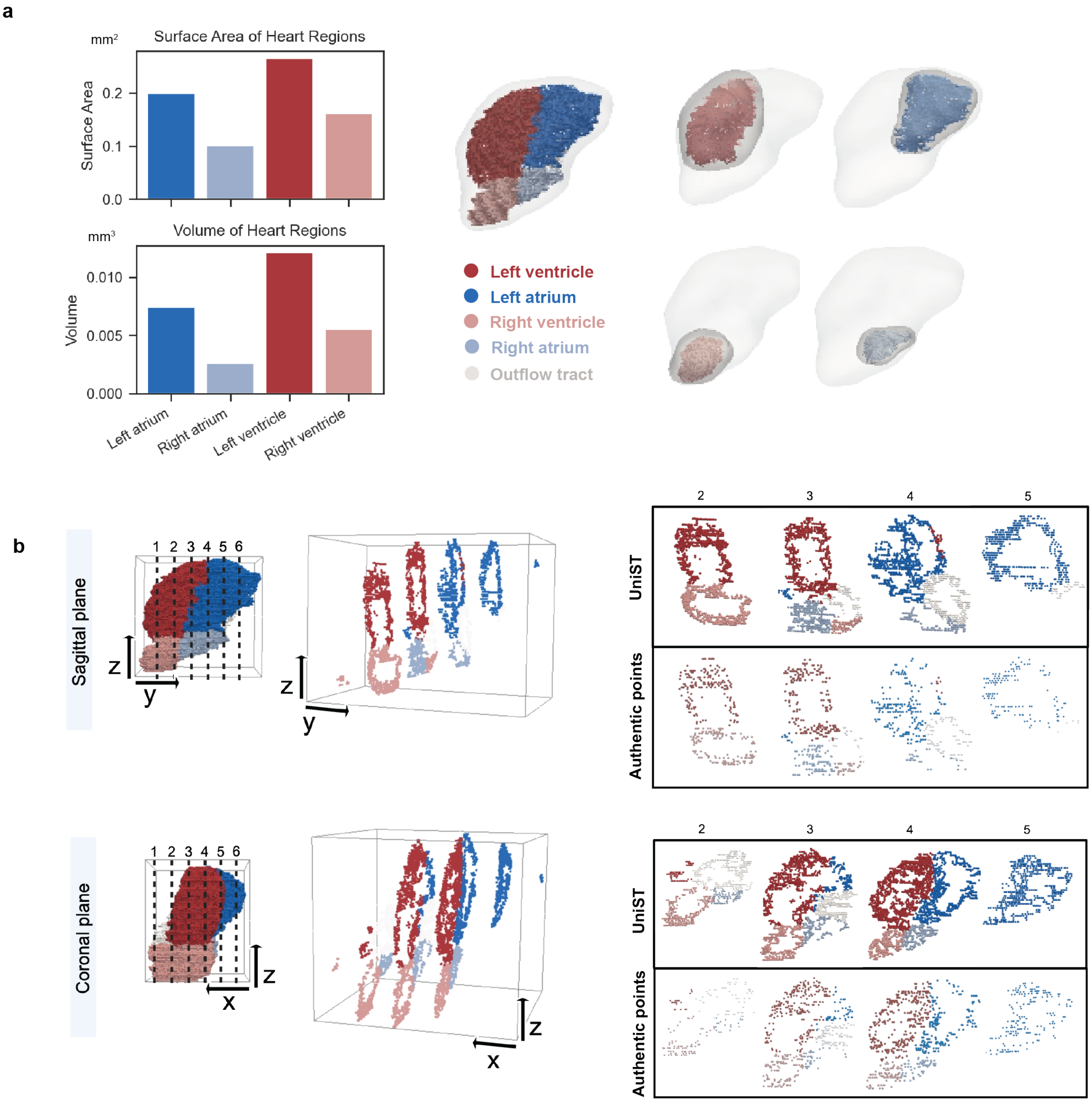

(a) Mesh reconstruction of heart regions with quantification of surface area and volume. (b) Pseudo-slice generation along the sagittal and coronal planes.

#### 2 List of Figures for Open-ST metastatic lymph node data

##### 2.1 Supplementary Figure 6. Preprocessing of Open-ST data to correct acquisition-induced cracks.

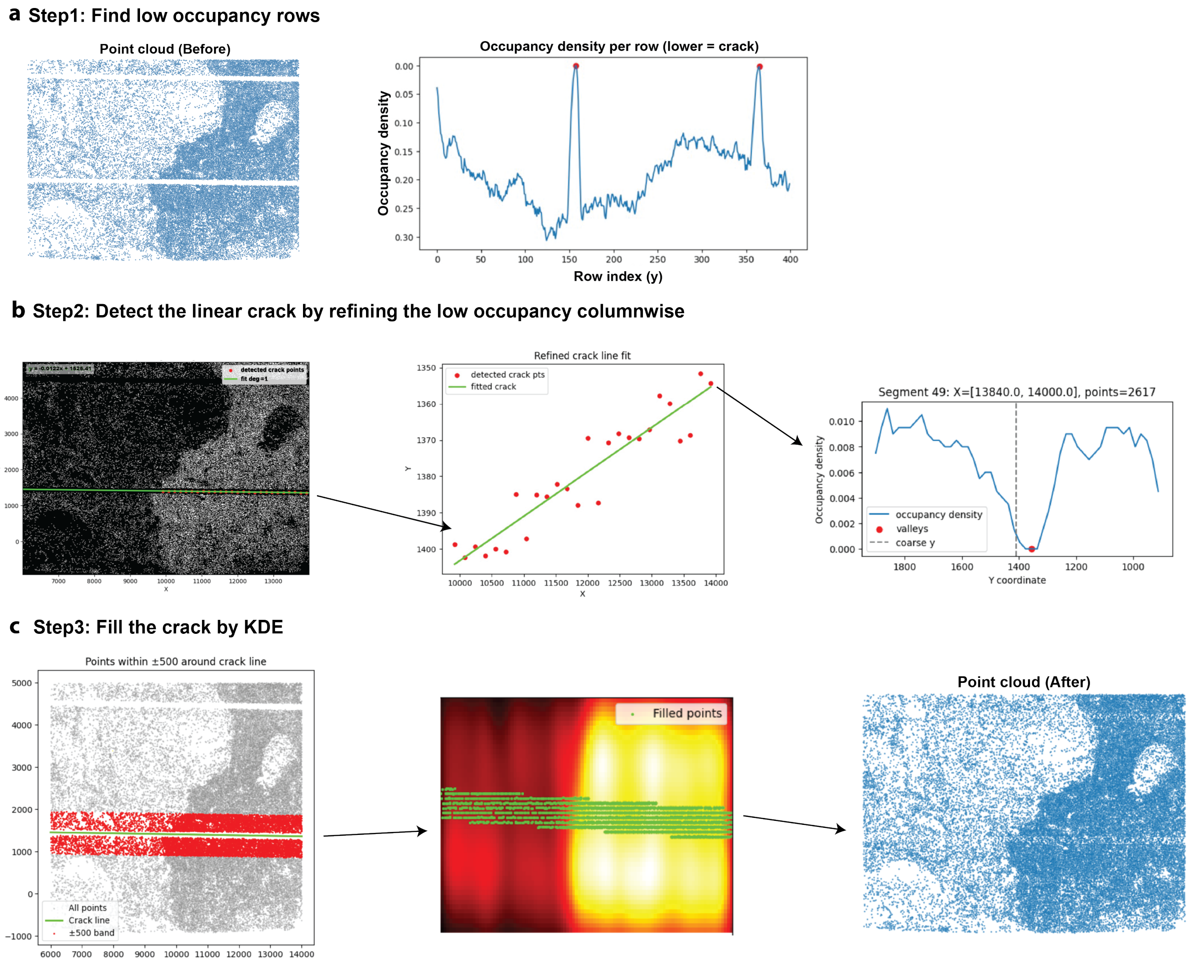

(a) Step1: After gridding ST slice into an occupancy image, horizontal rows with abnormally low occupancy were identified to detect candidate crack regions. (b) Step2: Bands surrounding the low-occupancy rows were extracted and low-occupancy grids within the band were then refined in a column-wise manner to delineate the linear cracks. (c) Step3: Detected cracks were filled using kernel density estimation (KDE)–based sampling to restore local point density, resulting in a continuous point cloud suitable for downstream reconstruction and analysis.

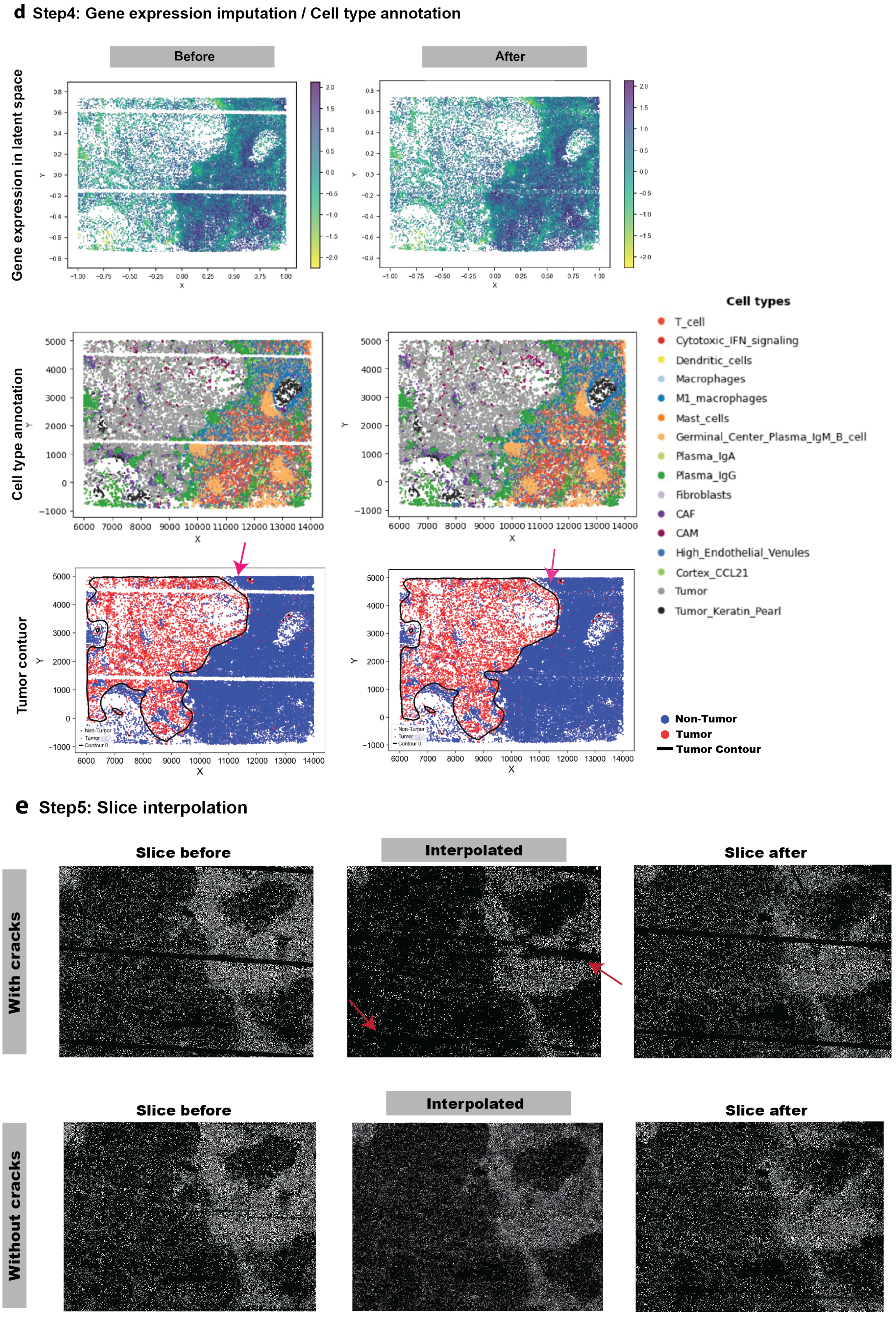

(d) Step4: Gene expression imputation, cell-type annotation and tumor-immune boundaries before and after cracks filled. (e) Step5: Slice interpolation results before and after cracks filled.

#### 2.2 Supplementary Figure 7. Visualization of reconstruction results.

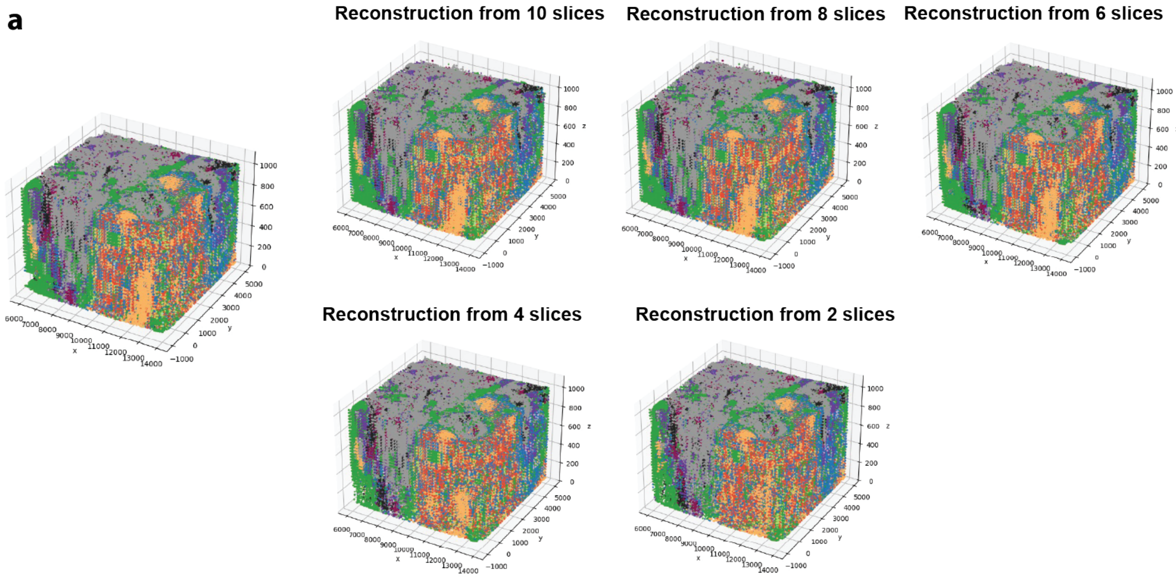

(a) 3D reconstruction results using 10, 8, 6, 4, and 2 slices. For visualization purposes, the z axis is scaled.

#### 2.3 Supplementary Figure 8. Reconstruction accuracy of tumor and T-cell regions.

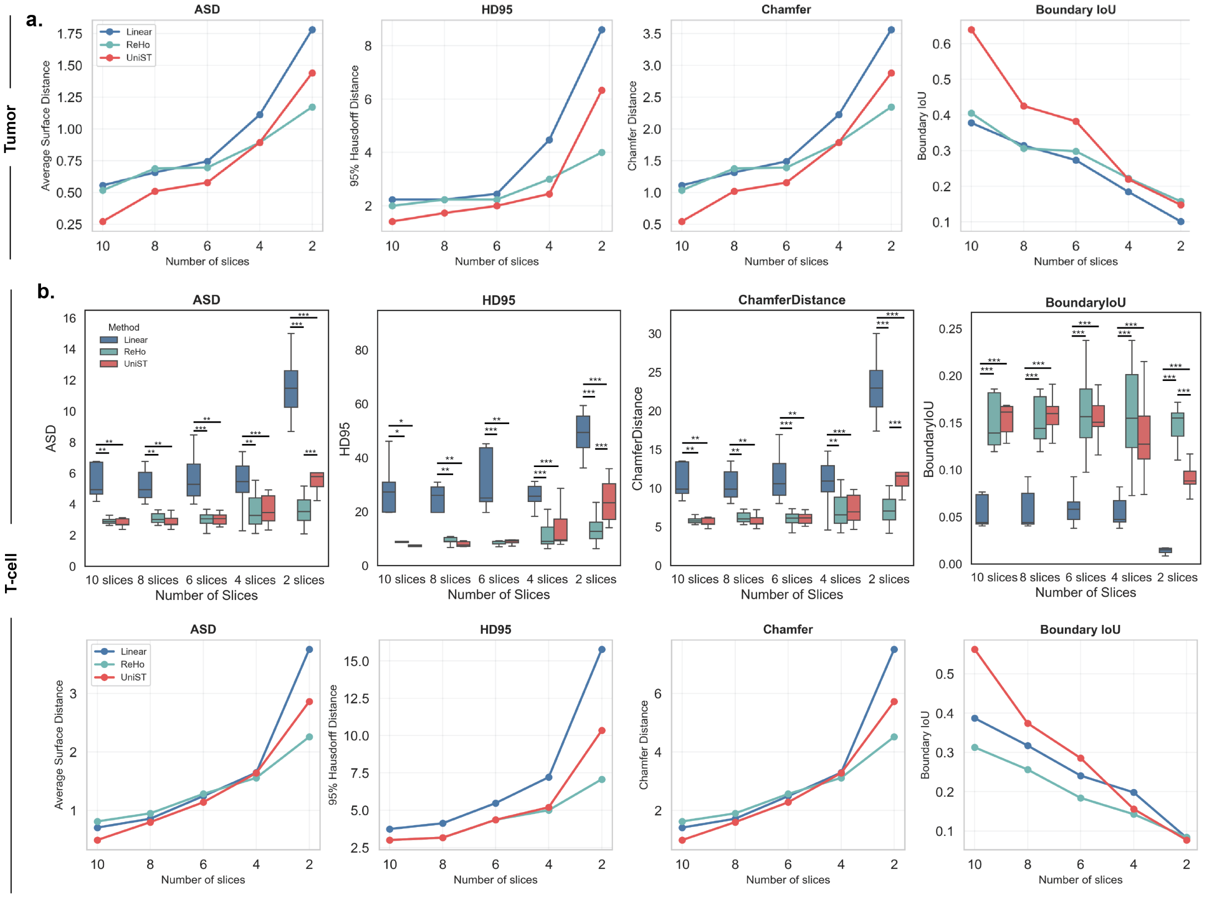

(a) ASD, HD95, Chamfer Distance and Boundary IoU computed at 3D level for reconstruction accuracy of the tumor region using 10, 8, 6, 4, 2 slices. (b) ASD, HD95, Chamfer Distance and Boundary IoU computed at both 2D (upper) and 3D (lower) levels for reconstruction accuracy of the T-cell region using 10, 8, 6, 4, 2 slices.

#### 2.4 Supplementary Figure 9. Reconstruction accuracy across 8 major tissue regions.

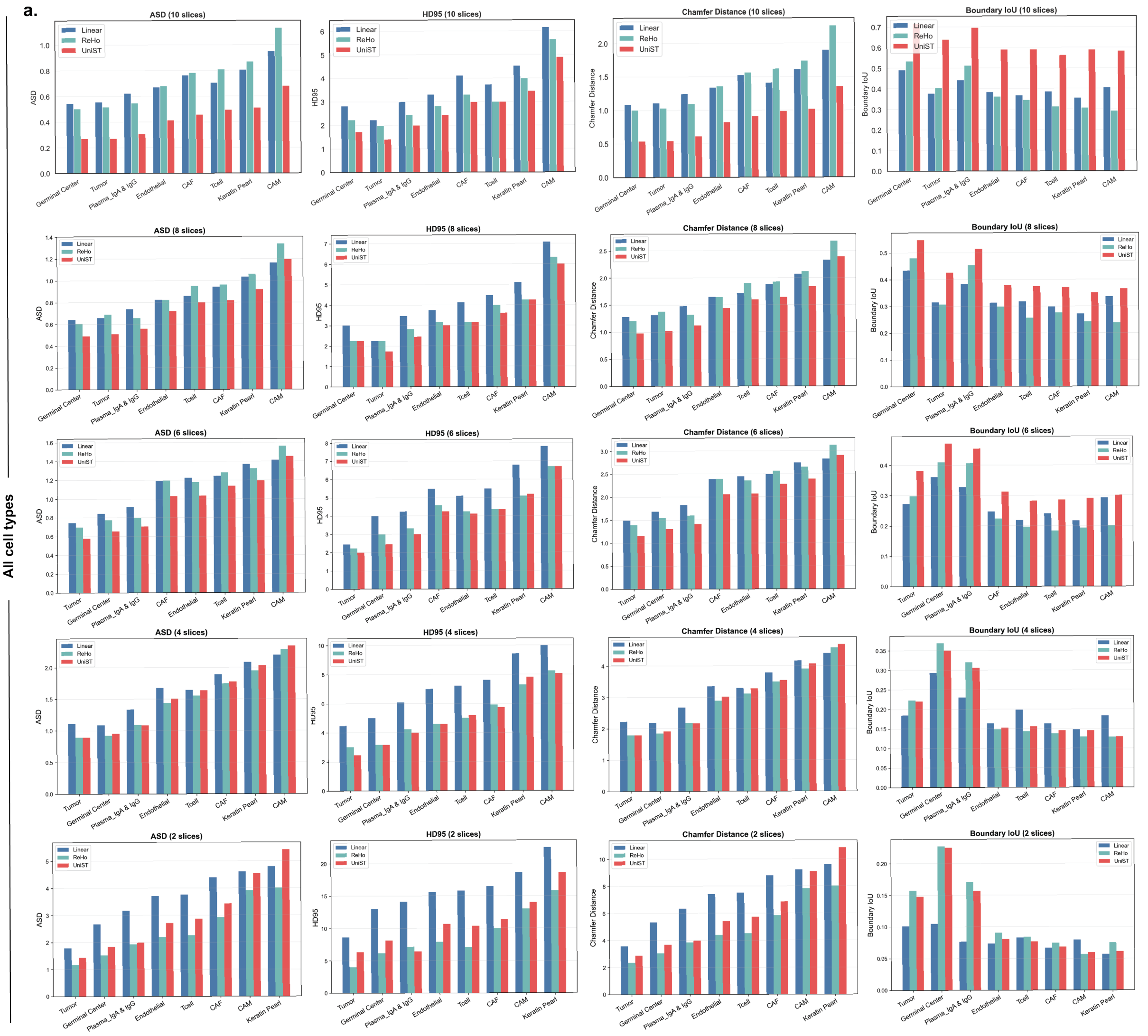

(a) ASD, HD95, Chamfer Distance and Boundary IoU computed at 3D level using 10, 8, 6, 4, 2 slices across 8 major tissue regions by three interpolation methods. Regions are ordered from left to right according to increasing ASD values obtained by UniST.

#### 2.5 Supplementary Figure 10. Gene expression distribution before and after UniST imputation.

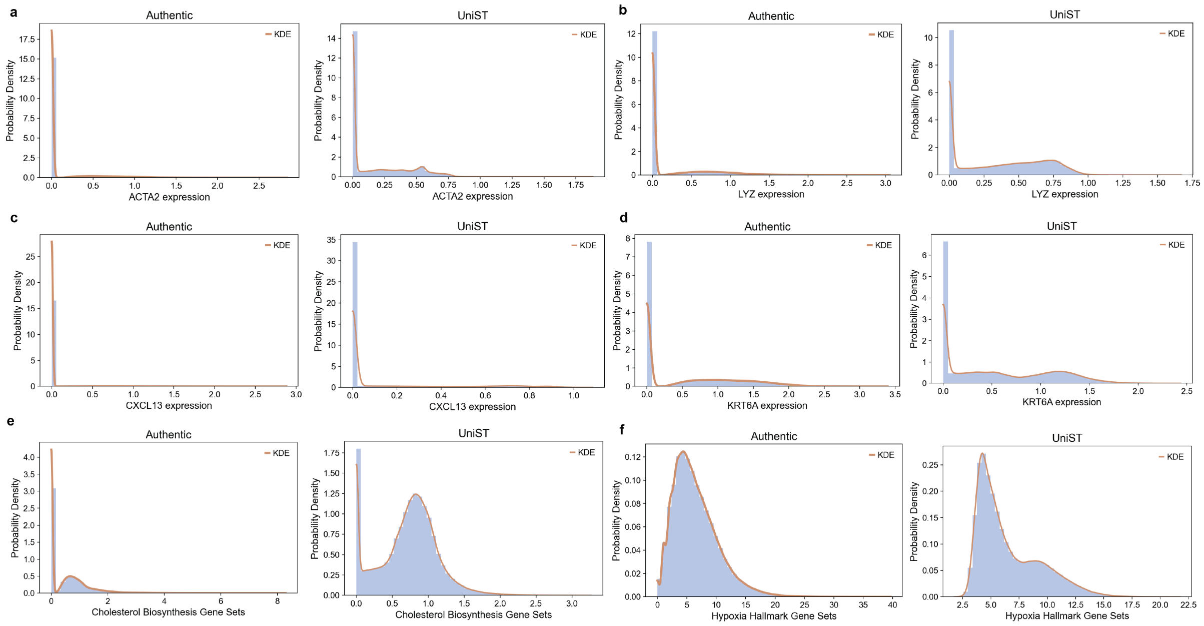

(a-d) Histograms with kernel density estimates (KDE) for the distribution of expression values for ACTA2 (a), LYZ (b), CXCL13 (c) and KRT6A (d) in the authentic data (left) and after UniST-based imputation (right). (e-f) Histograms with kernel density estimates (KDE) for the distribution of pathway enrichment scores for cholesterol biosynthesis gene sets (e) and hypoxia hallmark gene sets (f). Histograms were computed using 50 equally spaced bins.

### 3 List of Figures for Singular Genomics gastric carcinoma data

#### 3.1 Supplementary Figure 11. Preprocessing of gastric carcinoma tissues to patch the localized tissue loss.

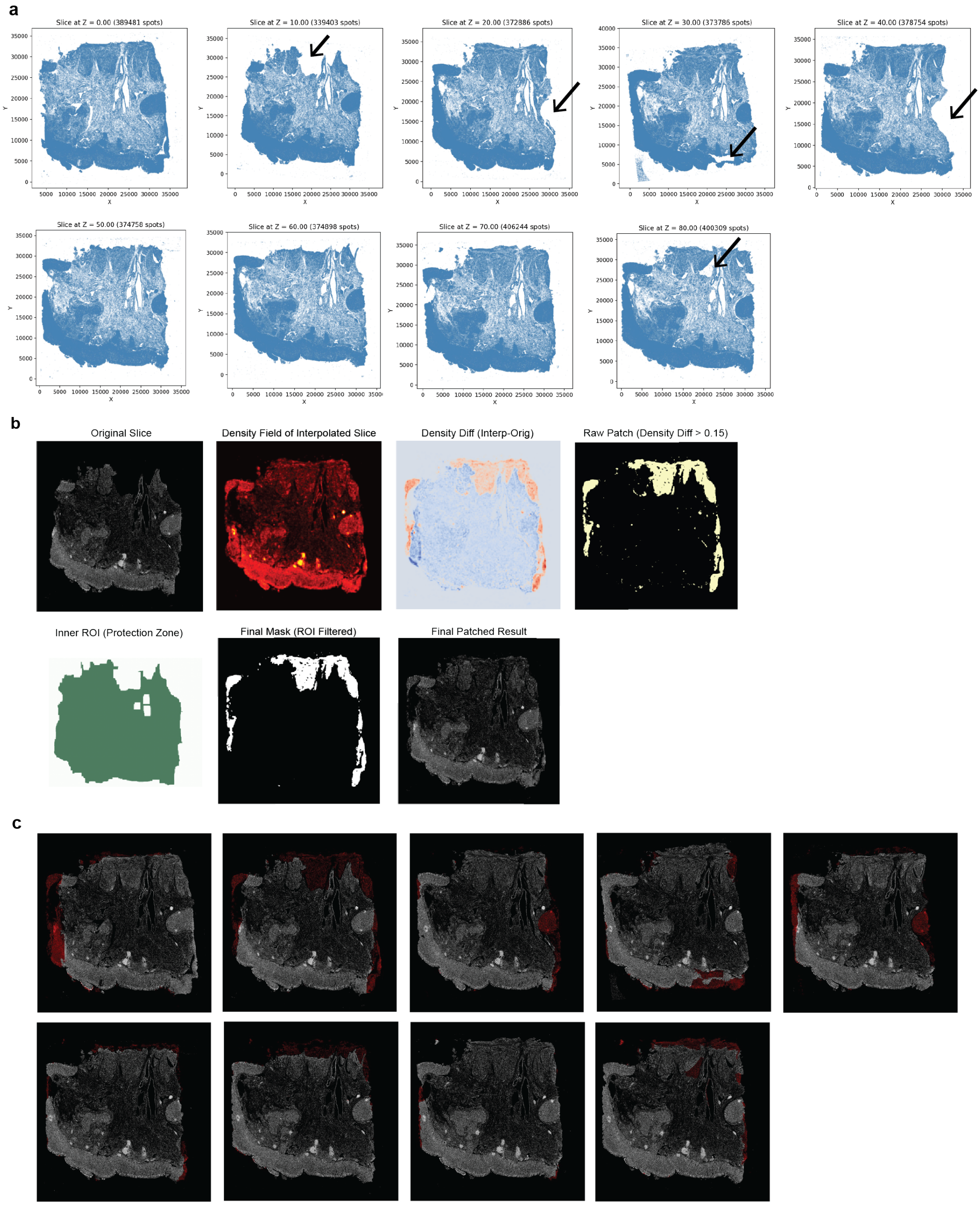

(a) Illustration of slices with tissue loss caused by acquisition-related artifacts. (b) The slice with tissue loss was patched using information interpolated from its immediately adjacent slices. To preserve original slice information, only the difference between the interpolated slice and the slice with tissue loss was patched, and was restricted to regions affected by tissue loss. (c) Patched results with patched regions highlighted in red color.

##### 3.2 Supplementary Figure 12. 3D reconstruction results.

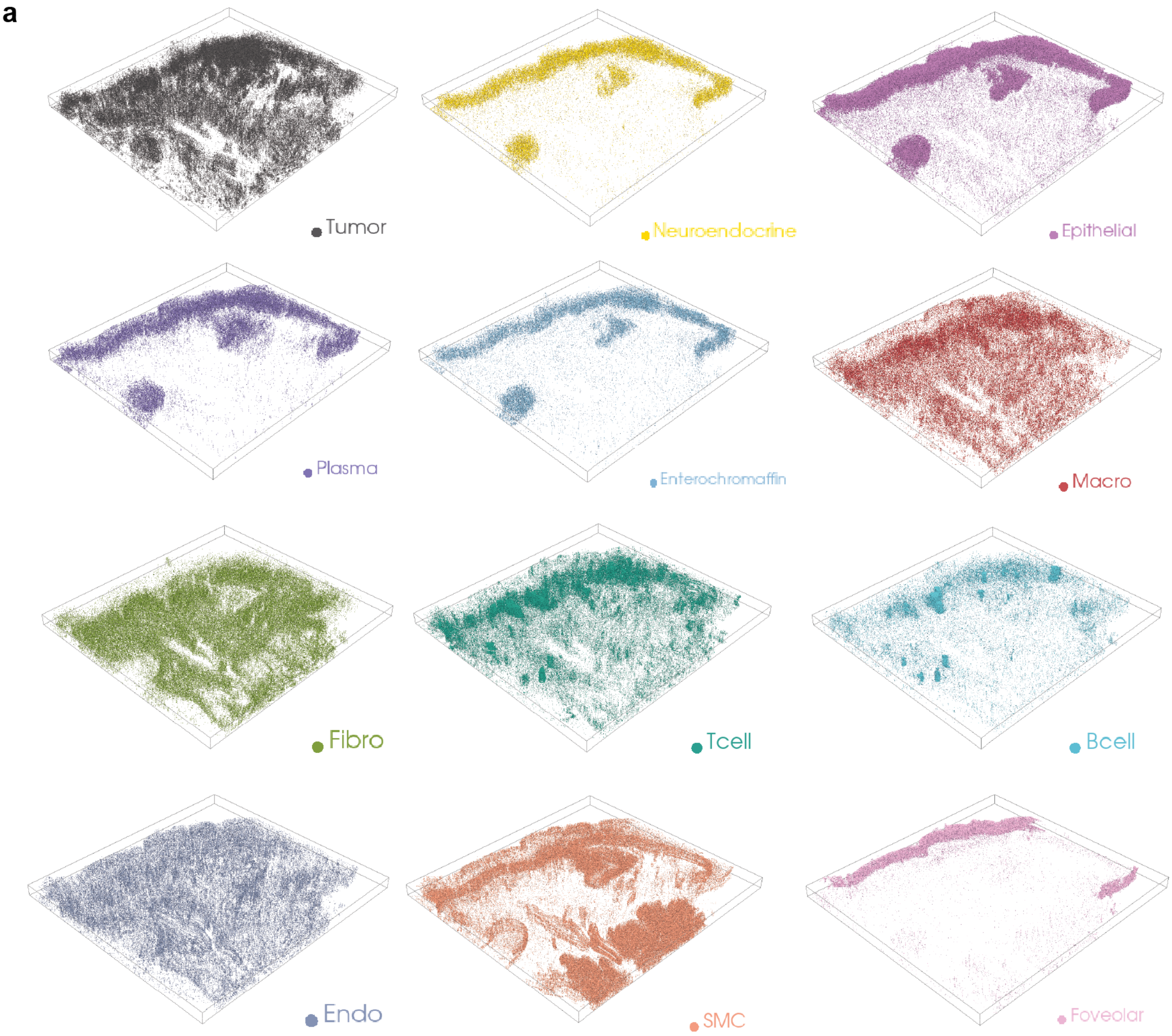

(a) 3D reconstruction results of 12 cell types.

#### 3.3 Supplementary Figure 13. Reconstruction accuracy across multiple regions and cell type composition

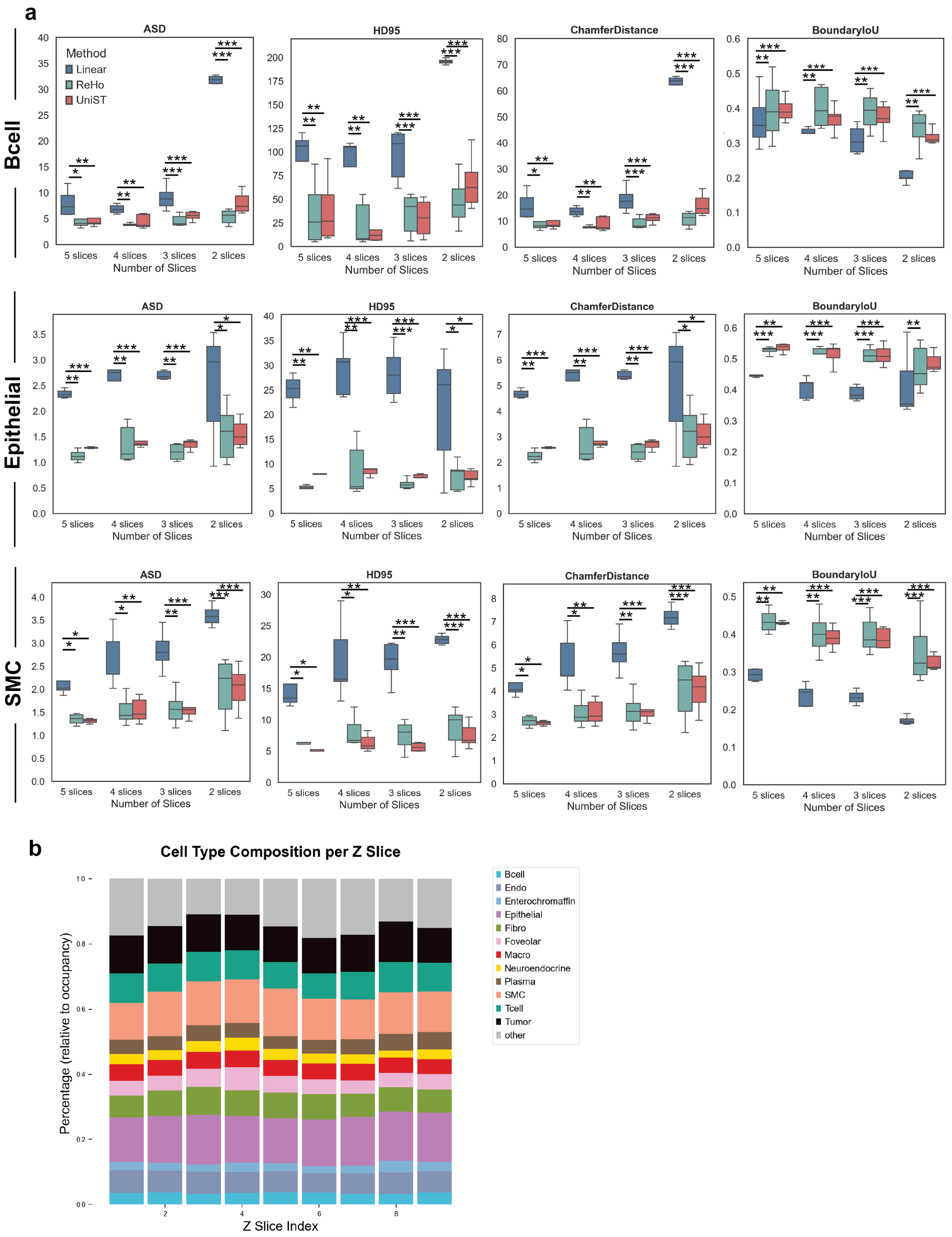

(a) ASD, HD95, Chamfer Distance and Boundary IoU computed at 2D level using 2-5 slices for three tissue regions. (b) Cell-type composition of major cell types across 9 slices.

##### 3.4 Supplementary Figure 14. Gene expression distribution before and after UniST imputation.

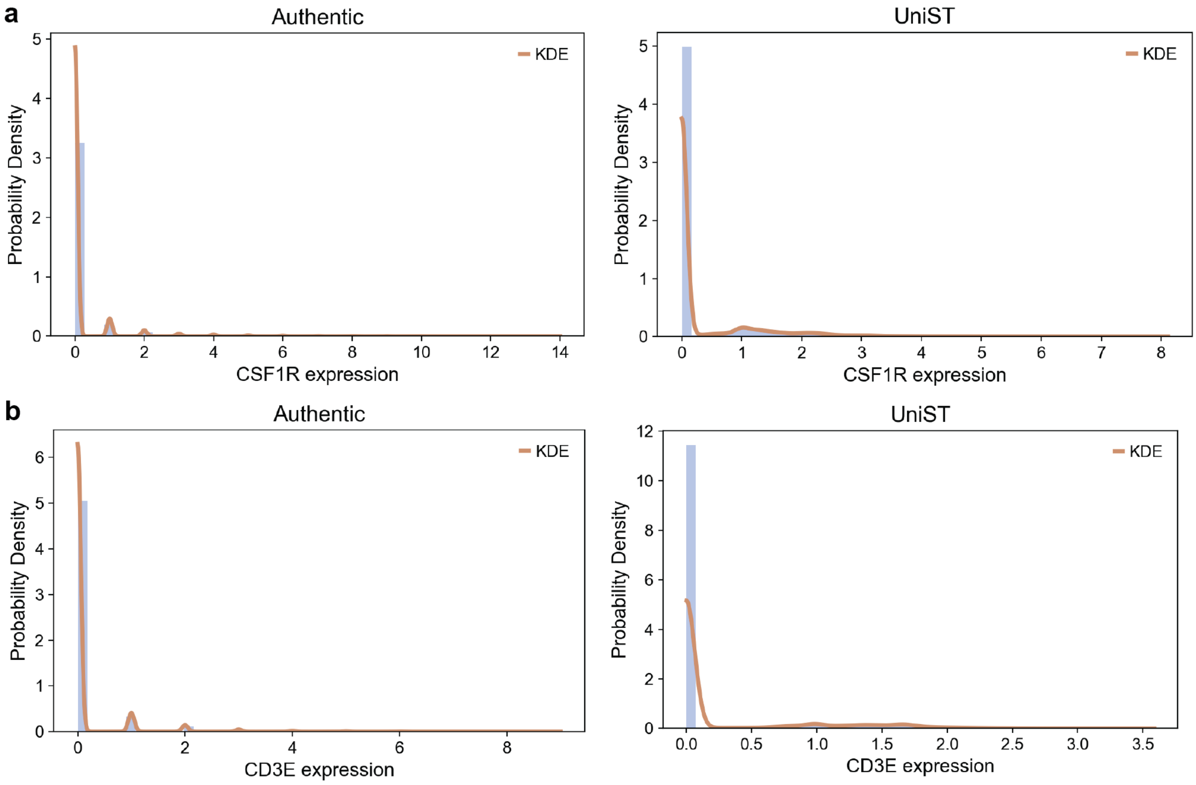

Histograms with kernel density estimates (KDE) for the distribution of expression values for CSF1R (a), CD3E (b) in the authentic data (left) and after UniST-based imputation (right). Histograms were computed using 50 equally spaced bins.

### Supplementary Notes

#### 4 Supplementary Note 1. Training loss functions.

##### 4.1 Point cloud upsampling

For the point cloud upsampling module, the training objective combines three complementary losses:

$$\mathcal{L} = \mathcal{L}_{\text{cd}} + 0.1\mathcal{L}_{\text{fit}} + 0.1\mathcal{L}_{\text{rep}}. \tag{1}$$

**(1) Chamfer Distance Loss:** Let the ground truth point cloud being $P_{gt} \in \mathbb{R}^{N_g \times 2}$,

$$\mathcal{L}_{\text{cd}} = \frac{1}{|P_u|} \sum_{p \in P_u} \min_{q \in P_{gt}} \|p - q\|_2^2 + \frac{1}{|P_{gt}|} \sum_{q \in P_{gt}} \min_{p \in P_u} \|q - p\|_2^2. \tag{2}$$

**(2) Fitting Loss:**

$$\mathcal{L}_{\text{fit}} = \frac{1}{N} \sum_{p \in P} \sum_{p_k \in P_k} \min_{p_r \in P_r} \left( \frac{\|p_k - (p_n - p)\|_2}{\sigma} \right)^2, \tag{3}$$

where $P$ denotes the set of input points, $P_k$ the kernel points, and $P_n$ the set of neighboring points within a ball query of radius $\sigma$.

**(3) Repulsive Loss:**

$$\mathcal{L}_{\text{rep}} = \frac{1}{N \times N_k \times (N_k - 1)} \sum_{p \in P} \sum_{p_k^i \in P_k} \sum_{i \neq j} \max\left(0,\, 1 - \frac{\|p_k^i - p_k^j\|_2}{\sigma}\right)^2. \tag{4}$$

##### 4.2 Slice interpolation

For the slice interpolation module, the training loss combines three terms:

$$\mathcal{L} \;=\; \lambda_1 \mathcal{L}_1 \;+\; \lambda_{\text{perc}} \mathcal{L}_{\text{perc}} \;+\; \lambda_{\text{style}} \mathcal{L}_{\text{style}}. \tag{5}$$

**(1) Reconstruction loss:**

$$\mathcal{L}_1 \;=\; \|\widehat{\mathbf{I}_\mathbf{t}} - \mathbf{I}_\mathbf{t}\|_1, \tag{6}$$

where $\mathbf{I}_\mathbf{t}$ is the ground-truth slice and $\widehat{\mathbf{I}_\mathbf{t}}$ is the interpolated slice.

**(2) Perceptual loss:** Let $\Psi_l(\cdot)$ denote the feature map at layer $l$ of the VGG-19.

$$\mathcal{L}_{\text{perc}} = \frac{1}{L} \sum_{l=1}^{L} \alpha_l \left\| \Psi_l(\widehat{\mathbf{I}_\mathbf{t}}) - \Psi_l(\mathbf{I}_\mathbf{t}) \right\|_1, \tag{7}$$

where $L$ is the number of layers used, and $\alpha_l$ is the weighting coefficient for that layer.

**(3) Style loss:**

$$\mathcal{L}_{\text{style}} = \frac{1}{L} \sum_{l=1}^{L} \alpha_l \left\| \mathbf{M}_l(\widehat{\mathbf{I}_\mathbf{t}}) - \mathbf{M}_l(\mathbf{I}_\mathbf{t}) \right\|_2, \tag{8}$$

where the Gram matrix $\mathbf{M}_l(X)$ is defined as

$$\mathbf{M}_l(X) = \Psi_l(X)^\top \Psi_l(X), \tag{9}$$

which quantifies the covariance of the VGG-19 features.

#### 4.3 Gene expression imputation

For the gene expression imputation module, we mentioned several losses:

**(1) The loss function to train GAE is masked Mean Square Error**, defined as:

$$\mathcal{L}_{\text{GAE}} = \frac{1}{|\mathbf{M}_y^+|} \sum_{i \in \mathbf{M}_y^+} \left(\hat{y}_i - y_i^{gt}\right)^2, \tag{10}$$

where $\mathbf{M}_y^+$ denotes the binary mask of $\mathbf{y}_{gt}$, and $\hat{\mathbf{y}}$ is the reconstructed value.

**(2) The loss function to train INR is also masked Mean Square Error**, defined as:

$$\mathcal{L}_{\text{embd}} = \frac{1}{|\mathbf{M}_z^+|} \sum_{i \in \mathbf{M}_z^+} \left(\hat{z}_i - z_i^{gt}\right)^2, \tag{11}$$

where $\mathbf{M}_z^+$ denotes the binary mask of $\mathbf{z}_{gt}$, and $\hat{\mathbf{z}}$ is the reconstructed latent representation.

**(3) The loss function for fine-tuning GAE decoder is defined as the combination of masked MSE, mean absolute error (MAE), and a Dice loss term**:

$$\mathcal{L}_{\text{recons}} = \frac{1}{|\mathbf{M}_y^+|} \sum_{i \in \mathbf{M}_y^+} (\hat{y}_i - y_i^{gt})^2 + \frac{1}{|\mathbf{M}_y|} \sum_{i \in \mathbf{M}_y} \left|\hat{y}_i - y_i^{gt}\right| + \lambda \, \mathbf{L}_{\text{dice}}, \tag{12}$$

where $\mathbf{M}_y^+$ denotes the set of non-zero entries, $\mathbf{M}_y$ is the full data inputs.

### 5 Supplementary Note 2. Model architectures and hyperparameters for gene expression imputation.

| **Component** | **Parameter** | **Value** |
|---|---|---|
| **GAE** | Graph Construction | Anisotropic kNN ($k = 6, w_z = 0.5$) |
| | Encoder Architecture | MLP (2048, 512, 64) |
| | Latent Dimension | 32 |
| | Learning Rate | $1 \times 10^{-5}$ (200 epochs) |
| **INR (FFN)** | Frequency Sampling ($\sigma_{x,y}$) | $\{1, 10, 100\}$ (Multi-scale) |
| | Frequency Sampling ($\sigma_z$) | $\{0.1, 1, 10\}$ (Multi-scale) |
| | Mapping Size ($m$) | 256 |
| | Hidden Layers | 2048 $\times$ 3 |
| | Learning Rate | $1 \times 10^{-4}$ (1000 epochs) |
| **Fine-tuning GAE** | Learning Rate | $1 \times 10^{-4}$ (1000 epochs) |
